# A Versatile Toolbox for Nanoscale Interrogation of Multiprotein Assemblies inside Living Cells

**DOI:** 10.1101/2025.04.30.651189

**Authors:** Arthur Felker, Michael Philippi, Michael Holtmannspötter, Christoph Drees, Evelin Schäfer, Martin Steinhart, Rainer Kurre, Changjiang You, Jacob Piehler

## Abstract

Quantitative analysis of protein interactions and the formation of higher-order assemblies in living cells remains a major challenge. Here, we introduce a versatile nanopatterning toolbox that employs capillary nanostamping of functionalized polymers to generate high contrast bio-functionalized nanodot arrays (bNDAs) with diameters below 500 nm. By leveraging orthogonal adaptor designs, we achieve robust immobilization of diverse fluorescent protein fusions, enabling simultaneous and selective recruitment of cytosolic and membrane-associated proteins into discrete nanodomains. This approach of forming cytosolic nanodot arrays (cNDAs) provides striking capabilities for dissecting cytosolic multiprotein complexes with molecular precision. Focusing on the assembly of the multimeric myddosome complex, we demonstrate density-dependent recruitment and co-localization of the core components MyD88, IRAK4, IRAK1, and TRAF6 within cNDAs. Super-resolution microscopy reveals distinct nanoscale clustering of MyD88 and IRAK4 and uncovers the ultrastructural architecture of IRAK4 oligomers. These analyses highlight the spatial organization and hierarchical assembly of the myddosome at the nanoscale in the native cellular context. Collectively, our findings establish cNDAs as a powerful platform for reconstituting and analyzing intricate multiprotein assemblies in live cells, offering new opportunities for elucidating the principles of complex protein networks.

## Introduction

Proteins are the core building blocks of life, orchestrating almost all cellular processes based on their intriguing variety of molecular functionalities. The huge complexity of protein functions can be explained by the association of proteins into complex assemblies via specific, noncovalent protein-protein interactions (PPIs). PPIs are often fast and transient, and therefore can be spatiotemporally regulated, e.g. via post-translational modifications, cofactors or interaction with other biomolecules.^1–4^ The resulting, intricate complex network of spatially and temporally regulated PPIs defines cellular identity and function, but also malfunction and disease, with a predicted average of 16 interaction partners for each protein.^5^ Despite tremendous progress in the structural prediction of protein complexes, quantitative interrogation of protein assemblies in the native cellular context remains a fundamental challenge in bioanalytics. Current quantitative PPI analysis is largely based on isolated proteins, making interaction analysis of multiprotein complexes highly demanding.^6, 7^ Furthermore, the *in vitro* assays lack the cellular context, e.g. localization at membranes, which can provide specific cues such as protein-lipid interactions and/or cooperative phenomena such as aggregation or condensation.^8, 9^

Surface micro- and nanopatterning techniques have emerged as powerful tools for systematically quantifying PPIs and protein-lipid interactions in live cells.^10–21^ While originally focusing on interactions at transmembrane receptors, the compatibility of these techniques for analyzing cytosolic PPIs has been recognized.^22–29^ However, spatial control and quantitative analysis of cooperative multiprotein interactions in the cellular context has remained highly challenging, requiring control of protein densities and composition. Recently, we introduced capillary nanostamping (CNS) for generating biofunctional nanodot arrays (bNDAs) suitable for capturing signaling complexes in living cells.^30, 31^ This approach combines top-down nanofabrication by CNS with bottom-up surface functionalization, leveraging fast and specific biomolecular recognition at the plasma membrane. Here, we have fundamentally advanced this technology to enable interrogating the assembly and dynamics of multiprotein complexes in the cytosol of live cells. To this end, we devised a set of carrier polymers with orthogonal functionalities for efficient CNS. In combination with optimized surface silanization, robust fabrication of bNDAs with < 500 nm diameter and 900 nm spacing was achieved (**Figure 1**A). This enabled efficient, high-density immobilization of target proteins. Versatility and capturing efficiency were further enhanced through engineering bifunctional adaptor proteins with orthogonal binding specificities. Expanding this approach to intracellular applications, we developed transmembrane adaptors enabling robust recruitment of cytosolic proteins into cytosolic nanodot arrays (cNDAs) (**Figure 1**B, C). We exploited these tools to spatially control the assembly of myddosomes, multiprotein complexes involved in relaying immune responses and inflammatory signals.^32, 33^ Myddosomes represent helical oligomeric assemblies comprising MyD88 and interleukin 1-associated kinases (IRAKs), which interact via their death domains to form stable, stacked structures.^34, 35^ By capturing MyD88 into cNDAs, we effectively induced MyD88 oligomerization, thereby facilitating the subsequent recruitment of downstream kinases. Combining our approach with DNA-PAINT imaging techniques^36^ allowed us to visualize the oligomeric nature of these complexes with molecular resolution in their cellular context. This system provides an exciting opportunity to achieve precise spatial control over intracellular proteins, enabling functional and structural interrogation of complex multiprotein assemblies in living cells.

**Figure 1.**
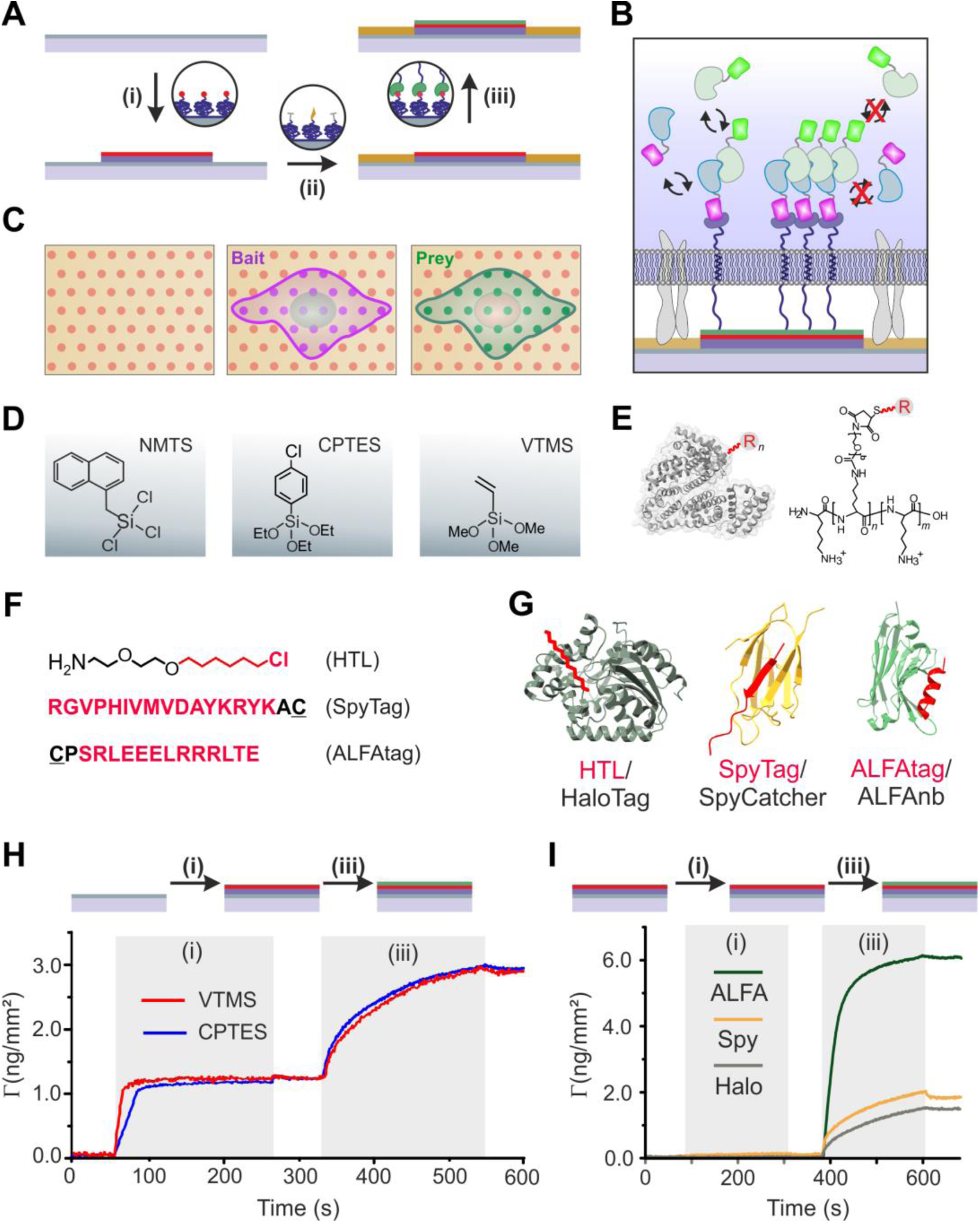
Concepts, reagents, and biomolecules employed for cNDA fabrication. (A-C) Cartoons depicting the fabrication and application of cytosolic NDAs for probing multi-protein complex formation inside live cells. (A) Preparation of bNDAs: (i) CNS of functionalized carrier biopolymers; (ii) Backfilling to inhibit nonspecific binding outside nanodots while ensuring cell adhesion; (iii) Biofunctionalization of nanodots with engineered adaptors to promote homomeric protein interactions. (B) Concept of cNDAs: Nanoscale enrichment of a transmembrane adaptor in high-density enables capturing cytosolic proteins via fluorescent protein tags (cyclic arrow below). Cooperative interactions within multiprotein complexes ensure formation and interrogation of multiprotein complex assembly inside cells (blocked cyclic arrows). (C) Top view of: bNDAs, and cNDAs of bait and prey (left to right). (D) Silanes used for surface silanization. (E) Functionalized carrier polymers BSA (left) and PLL-PEG (right), with the functional group indicated as ‘R’. (F, G) Functional groups used for surface biofunctionalization (F) and corresponding binding proteins for fusion (G): HaloTag (Halo, PDB:6U32), SpyCatcher (SpyC, PDB:4MLI) and ALFAtag-Nanobody (ALFAnb, PDB:6I2G) in complex with their binding partner (highlighted in red). (H) Comparison of PPH (i) binding on surfaces silanized with CPTES and VTMS, respectively, followed by capturing of HaloTag-mEGFP (iii). (I) Surface loading for differently functionalized PLL-PEG derivatives. Injection of mEGFP as a negative control (i) followed by the corresponding binder fused to mEGFP (iii).

## Results and Discussion

### Tailored silanization yields effective surface functionalization with carrier polymers

We have previously established CNS of functionalized bovine serum albumin (BSA) on glass surfaces rendered hydrophobic by silanization with 1-naphthylmethyltrichlorosilane (NMTS) to achieve high contrast bNDAs.^31^ Here, we extended and broadly enhanced the capabilities of bNDAs as a generic approach by optimizing surface silanization, carrier polymers, and orthogonal protein recognitions (**Figure 1**D-G). To this end, we turned to functionalized poly-*L*-lysine graft poly(ethylene glycol) block copolymers (PP, **Figure 1**E), which ensures much improved reproducibility and quality control. PP furthermore enables powerful surface engineering of surface properties such as minimizing non-specific binding of proteins and other compounds involved in cell culturing and high-resolution microscopic assays.^37, 38^

However, surface coating by PP is based on electrostatic interactions with oxide surfaces such as silica. Therefore suitable silanizations yielding hydrophobicity and negative potential are required for their application in CNS. Due to the limited availability of NMTS, strong nonspecific binding of aromatic organic compounds, and limited negative potential, we explored two other silanes, 4-chlorophenyltriethoxysilane (CPTES) and vinyltrimethoxysilane (VTMS), as potential candidates (**Figure 1**D). Silanization of glass substrates with CPTES yielded a substantial increase in hydrophobicity (θ = 54.4±1.4°, Figure S1A). A considerably lower surface wetting was observed upon silanization with VTMS (θ = 82.3±1.2°), in line with the superhydrophobic characteristics reported for this silane.^39^ For surface functionalization by CNS, we synthesized PP conjugates with the HaloTag ligand^40^ (PPH), the ALFA-tag^41^ (PPA) and the SpyTag003^42^ (PPS), respectively (**Figure 1**F, G). We have previously successfully applied the highly bioorthogonal HaloTag/HTL reaction for surface micro- and nanopatterning using different carrier polymers.^16, 17, 31^ However, the reaction is relatively slow (∼10^3^-10^4^ M^-1^s^-1^),^43^ limiting capturing efficiencies at low concentrations of the target protein. We therefore explored suitability of the optimized SpyTag/SpyCatcher reaction, for which a much higher association rate constant (*k_on_* ≈ 5×10^5^ M^-1^s^-1^) has been reported, while still being irreversible due to the formation of an isopeptide bond.^42^ Recognition of the ALFAtag by the ALFAnb is similarly fast (∼4×10^5^ M^-1^s^-1^), but the tag is highly resistant to chemical modification such as fixation.^41^ A potential drawback is the reversibility of this interaction, though with a very low dissociation rate constant. As a reference, corresponding derivates of BSA with HTL and ALFAtag, respectively, were synthesized.

Binding kinetics of the carrier polymers on differently silanized surfaces was characterized by surface sensitive detection using reflectance interference and total internal reflection spectroscopy (RIF-TIRFS) in a flow-through system.^44^ Fast, irreversible binding of PP derivatives was observed on silanized glass substrates with overall binding characteristics similar to what we have previously observed for hydrophilic glass surfaces (**Figure 1**H, Figure S1B). Surprisingly, binding to VTMS-silanized glass was faster as compared to CPTES, reaching saturation within a few seconds. The maximum surface loading of PPH and subsequent binding of the interaction partner was largely independent of the type of silanization (**Figure 1**H, Figure S1C). However, PPA showed the overall highest binding capacity, which may be explained by higher reactivity and availability on surface (**Figure 1**I, Figure S1D).

### High contrast bNDAs are achieved by diverse functionalized carrier polymers

With this toolbox of functionalized carrier polymers in hand, we systematically explored their performance for fabricating bNDAs for efficient protein capturing. After CNS of PP derivatives on differently silanized substrates and backfilling with non-functionalized PPM (M stands for <u>M</u>ethoxy groups), the obtained bNDAs were stained by incubation of the corresponding binding protein fused to GFP (HaloTag-mEGFP, SpyCatcher-sfGFP and ALFAnb-mEGFP, respectively). Representative total internal reflection fluorescence (TIRF) microscopy images of bNDAs of PPS and PPA on CPTES and VTMS, respectively, are shown in **Figure 2**A-F. Diffraction-limited TIRF imaging confirmed highly selective and homogeneous GFP capturing into bNDAs for different combinations of substrate silanization and carrier polymer functionalization (**Figure 2**A-F, Figure S2). More detailed quantification revealed strong differences in the nanodot peak intensities for different conditions (**Figure 2**G). Notably, CNS performed on VTMS-silanized surfaces consistently yielded substantially higher nanodot peak intensities as compared to CPTES-silanized surfaces. Moreover, improved nanodot peak intensity was achieved for SpyTag/SpyCatcher as compared to HTL/HaloTag, which was also more pronounced on VTMS-silanized surfaces. However, ALFAtag-mediated capturing exhibited by far the best performance, yielding peak intensities up to 40-fold higher compared to HaloTag. These differences are in line with the different binding capacities observed by RIF-TIRFS detection, but also highlight that further improvement can be achieved by CNS on super-hydrophobic surfaces. Surprisingly, the enhanced nanodot binding capacities were only partially reflected in the resulting contrast (**Figure 2**H). Increased signals within nanodots correlated with elevated background fluorescence between nanodots, even though identical PPM backfilling was employed for minimizing nonspecific protein adsorption. Given the < 900 nm spacing between adjacent nanodots, we therefore interpreted the increased background signals were caused by optical cross-talk between diffraction-limited images of the nanodots, which was also observed previously.^31^

**Figure 2.**
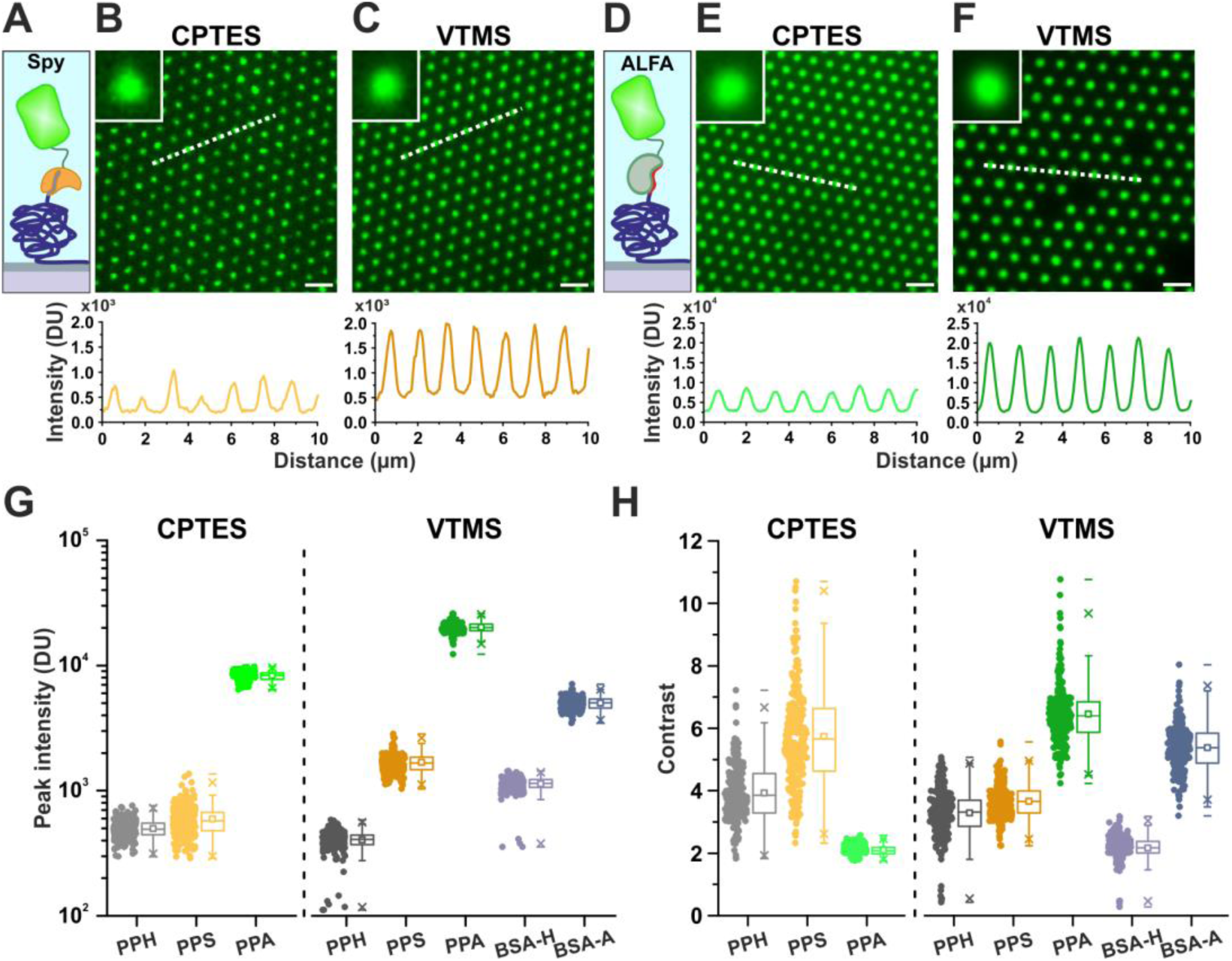
Orthogonal binding strategies for high-contrast bNDAs on silanized surfaces. (A-C) Schematic representation of the PPS/SpyCatcher-sfGFP biofunctionalization strategy (A), alongside TIRF microscopy images illustrating bNDAs on CPTES (B) and VTMS (C) surfaces. White dashed lines denote the regions selected for intensity profiles. Scale bars: 2 µm. Insets show enlarged views of individual nanodots, highlighting fluorescence distribution and uniformity. (D-F) Schematic overview of the PPA/ALFAnb-mEGFP biofunctionalization strategy (D), with corresponding TIRF microscopy images on CPTES (E) and VTMS (F) surfaces. White dashed lines denote regions used for intensity profiles. Scale bars: 2 µm. Insets show enlarged views of individual nanodots. (G) Quantitative analysis of fluorescence peak intensity values across different PP and BSA derivatives on various surfaces. Each dot in the box plot represents the value from a single nanodot. (H) Contrast analysis of bNDAs across different PP and BSA derivatives on different surfaces. Each dot in the box plot represents the value from a single nanodot.

### Ultrahigh contrast, sub-500 nm diameter bNDAs uncovered by DNA-PAINT imaging

In order to resolve the functional morphology of bNDAs at the nanoscale, we turned to super-resolution (SR) imaging by DNA-PAINT.^36^ Focusing on ALFA-tag functionalized NDAs, we implemented two distinct labeling approaches using nanobodies conjugated to suitable DNA docking strands, *i.e.*, anti-GFP nanobody (GFPnb) for two-step labeling (**Figure 3**A) and ALFAnb for direct labeling (Figure S3A). The two-step labeling strategy exploited the well-established staining of GFP fusion proteins via GFPnb.^45^ For this purpose, NDAs were firstly stained with GFP-tagged ALFAnb, as done for diffraction-limited imaging, and subsequently incubated with commercially available GFPnb conjugated with a docking strand (**Figure 3**A). For the direct labeling approach, NDAs were incubated with commercially available ALFAnb conjugated with a docking strand (Figure S3A). Representative SR images comparing PPA bNDAs on CPTES and VTMS by these two approaches are shown in **Figure 3**B-D and Figure S3B-D.

**Figure 3.**
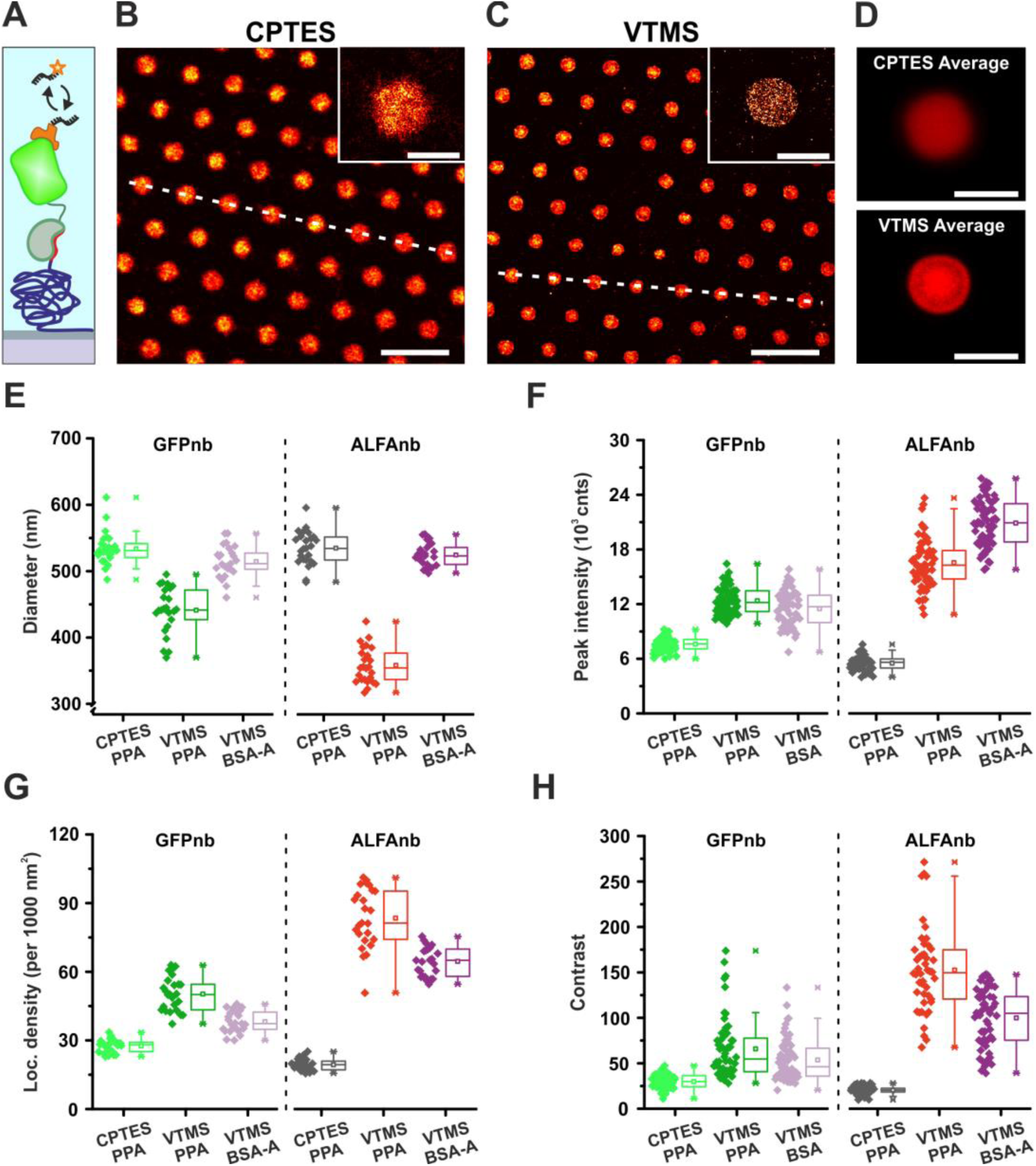
Nanoscale structural characterization of bNDAs using DNA-PAINT super-resolution imaging. (A) Schematic illustration of the DNA-PAINT labeling strategy applied to PPA/ALFAnb-mEGFP bNDAs, utilizing an anti-GFP nanobody for targeted imaging. (B, C) Representative DNA-PAINT super-resolution images on CPTES (B) and VTMS (C) surfaces. White dashed lines denote regions analyzed for intensity profiles shown in Figure S3. Scale bars: 2 µm. Insets provide magnified views of individual nanodots, revealing sub-diffraction-limit structural features. Scale bars: 500 nm. (D) Averaged DNA-PAINT image generated by aligning and averaging 200 individual nanodots. Scale bars: 500 nm. (E-H) Quantitative characterization of nanodots from DNA-PAINT super-resolution data comparing diameter (E), peak intensity (F), localization density per 1000 nm² (G), and contrast (H). Data was analyzed across different combinations of surface silanization methods, carrier bioconjugates, and imaging strategies (GFPnb, ALFAnb). Each data point in the boxplot represents an individual nanodot.

A typical localization precision of ∼6 nm was achieved in these experiments, thus enabling to clearly resolve the morphology and heterogeneity of individual nanodots. Significant differences were observed depending on the silanization chemistry employed, with bNDAs on VTMS-silanized surfaces exhibited smaller dot sizes, along with higher functionalization densities as compared to CPTES (**Figure 3**D-G). The much more well-defined contour of nanodots on VTMS-vs. CPTES-silanized surfaces could be clearly discerned upon averaging images of individual nanodots (**Figure 3**D), which can be explained by better liquid confinement during CNS due to lower substrate hydrophobicity.^46^ Control experiments using CNS of BSA-ALFA on VTMS-silanized surfaces confirmed substantial improvement in spatial confinement obtained by PLL-PEG (**Figure 3**E-H, Figure S4). Very similar results were obtained employing directly labeling by ALFAnb conjugated to the same docking strand (**Figure 3**E-G, Figure S3, Figure S4). Strikingly, coherent and much lower background levels, and correspondingly much higher contrasts were observed for super-resolved DNA-PAINT images, reaching values of up to 150 for PPA on VTMS (**Figure 3**H). This observation corroborates that spectral cross-talk from nanodots accounts for the high apparent background signals and the correspondingly low contrast values observed by diffraction-limited TIRF imaging (cf. **Figure 2**). Overall, super-resolution imaging of bNDAs highlights the substantially improved capabilities of functionalized PP printed on VTMS-silanized substrates, which was applied as the method of choice in the following experiments.

### Orthogonal adaptors enable binary control of target protein capturing into bNDAs

We have previously demonstrated that adaptor proteins can enhance binding capacity and capturing efficiency in the case of surface functionalization with relatively slow association kinetics such as HTL.^31^ Building on this concept, we here designed a set of orthogonal adaptor proteins to enable selective capturing via GFP- and mCherry-tagged proteins (**Figure 4**A, C, Figure S5A-I). To this end, we made use of corresponding designed ankyrin repeat proteins (DARPins), which are based on highly stable scaffolds and therefore maintain structural integrity after immobilization on surfaces.^47^ For binding GFP derivatives, we used the GFP clamp,^48^ and for mCherry the DARPin 2m22.^49^ Both proteins were used in tandem (tdR7, td2m22) manner and combined with different binders for targeting into bNDAs. Fast, efficient and highly specific capturing of mEGFP and mCherry by the corresponding adaptors bound to PP functionalized surfaces was verified by RIFS detection (Figure S5J, K). To evaluate the performance in bNDAs, adaptor proteins were incubated after CNS of correspondingly functionalized PP derivatives, followed by staining with 500 nM mEGFP and mCherry for 15 min, respectively. Typical bNDAs obtained under these conditions are shown in **Figure 4**B, D, and Figure S5A, B, E-G.

**Figure 4.**
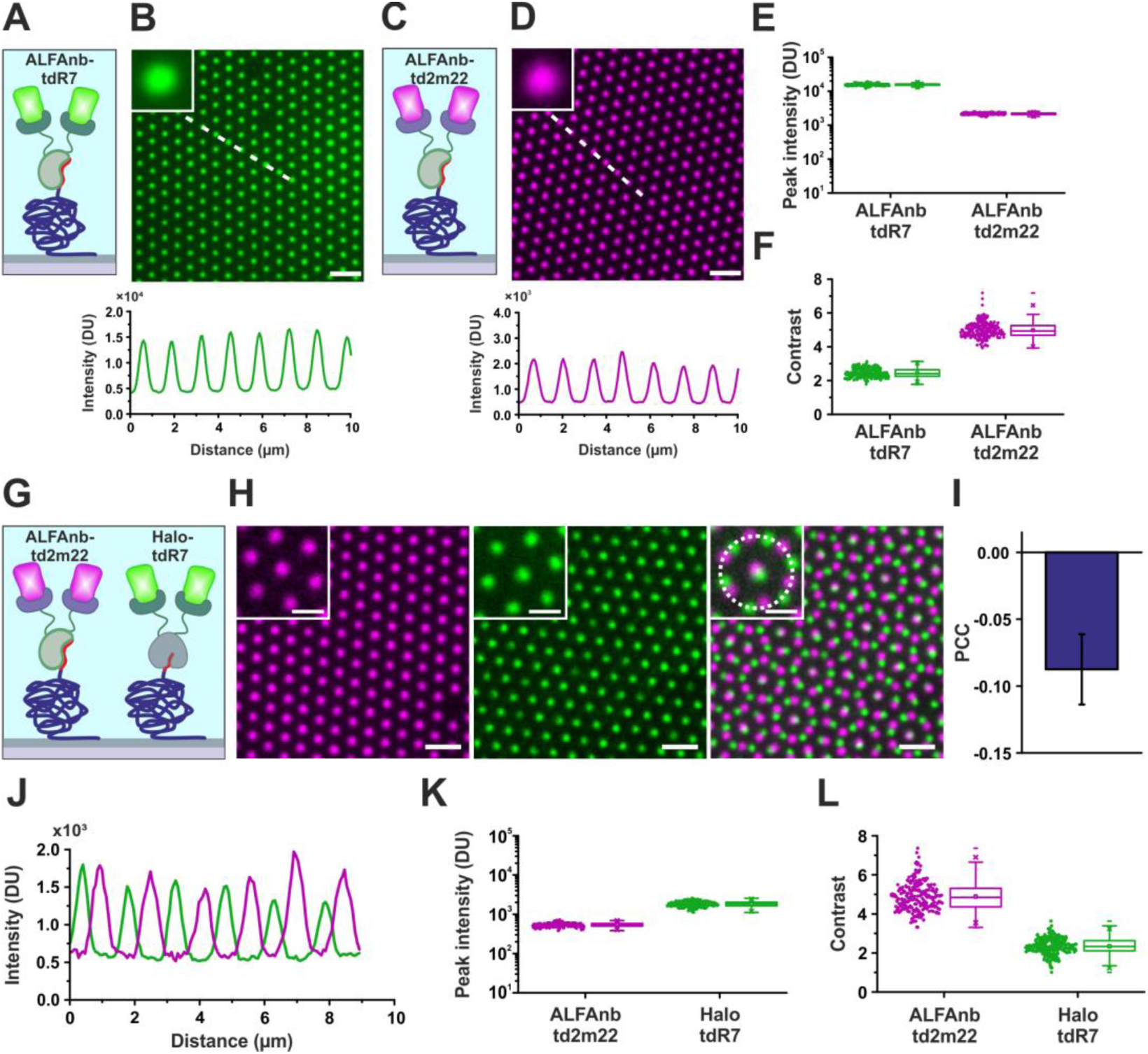
Bifunctional adaptor proteins for orthogonal capturing of fluorescent proteins. (A, C) Schematic representation of the molecular framework governing the selective immobilization of mEGFP (A) or mCherry (C) via PPA/ALFAnb-tdR7 or ALFAnb-td2m22 (B, D) Representative TIRF microscopy images of tdR7/mEGFP (B) and td2m22/mCherry (D). Insets show enlarged views of individual nanodots. White dashed lines indicate regions selected for intensity profiles. Scale bars: 2 µm. (E, F) Quantitative analysis of peak fluorescence intensity (E) and contrast (F) for tdR7/mEGFP and td2m22/mCherry. (G) Schematic illustration of the multiplexed biofunctionalization approach combining PPA/ALFAnb-td2m22 and PPH/Halo-tdR7. (H) Representative dual-color TIRF microscopy images showing spatially distinct capture (left, middle) and merged channels (right). Insets provide detailed views of individual channels and the composite merged image. Scale bars: 2 µm; insets: 1 µm. (I) PCC analysis quantifying co-localization within the binary patterning framework. (J) Fluorescence intensity line profiles corresponding to (H). (K, L) Quantitative analysis of fluorescence peak intensities (K) and contrast (L) for respective channels.

Very high peak intensities and contrasts were achieved, in line with the increased on-rate of the adaptor binders (**Figure 4**E, F, Figure S5). Leveraging capturing efficiency and specificity of these orthogonal adaptors, we generated more complex bNDAs by applying binary patterning approaches. To this end, PPH and PPA were subsequently stamped onto the same substrate, leading to two spatially shifted NDAs (**Figure 4**G). After functionalization with the corresponding adaptor proteins (ALFAnb-td2m22 and Halo-tdR7), co-incubation of mEGFP and mCherry resulted in two distinct bNDAs in the green and orange emission channels, exhibiting high peak intensity and contrast values (**Figure 4**H, K, L). Spatial co-localization analysis confirmed strong anticorrelation, demonstrating the capability of orthogonal protein targeting into spatially separated nanodots (**Figure 4**I, J). A similar level of anticorrelation was observed for sequential printing of PPA and PPS, and subsequent staining with ALFAnb-Dy647 and SpyCatcher-sfGFP, respectively (Figure S6A-F). Moreover, complementary, efficient binary bNDAs were achieved not only by functionalizing PPA-NDAs with two differently fluorescent-labeled binders (Figure S6G-L), but also by using a mixture of orthogonal adaptor proteins (ALFAnb-tdR7 and ALFAnb-td2m22) for capturing mEGFP and mCherry into the same bNDAs (Figure S6M-R). These results highlight the versatile opportunities opened by combining orthogonal surface functionalization with distinct orthogonal adaptor binders in NDAs for controlling spatial co-organization of proteins at different scales.

### Transmembrane adaptors for cytosolic NDAs of proteins in live cells

For nanopatterning of proteins inside live cells, we devised corresponding transmembrane adaptor proteins capable of recruiting cytosolic bait proteins into cNDAs at the plasma membrane. For stable and high-affinity capturing of GFP-tagged cytosolic proteins, the established GFPnb^45^ was C-terminally fused to an artificial transmembrane domain (TMD) and an extra-cellular ALFAnb (GFPnb-TA) as schematically depicted in **Figure 5**A. Efficient targeting of these transmembrane adaptors into the plasma membrane was ensured by an N-terminal signal peptide. For proof-of-concept experiments, HeLa cells co-expressing GFPnb-TA and mEGFP were cultured on PPA NDAs, which were backfilled with RGD-functionalized PP (PP-RGD) to promote cell adhesion.^37^ Live-cell TIRF imaging revealed efficient recruitment of cytosolic mEGFP into cNDAs (**Figure 5**B).

**Figure 5.**
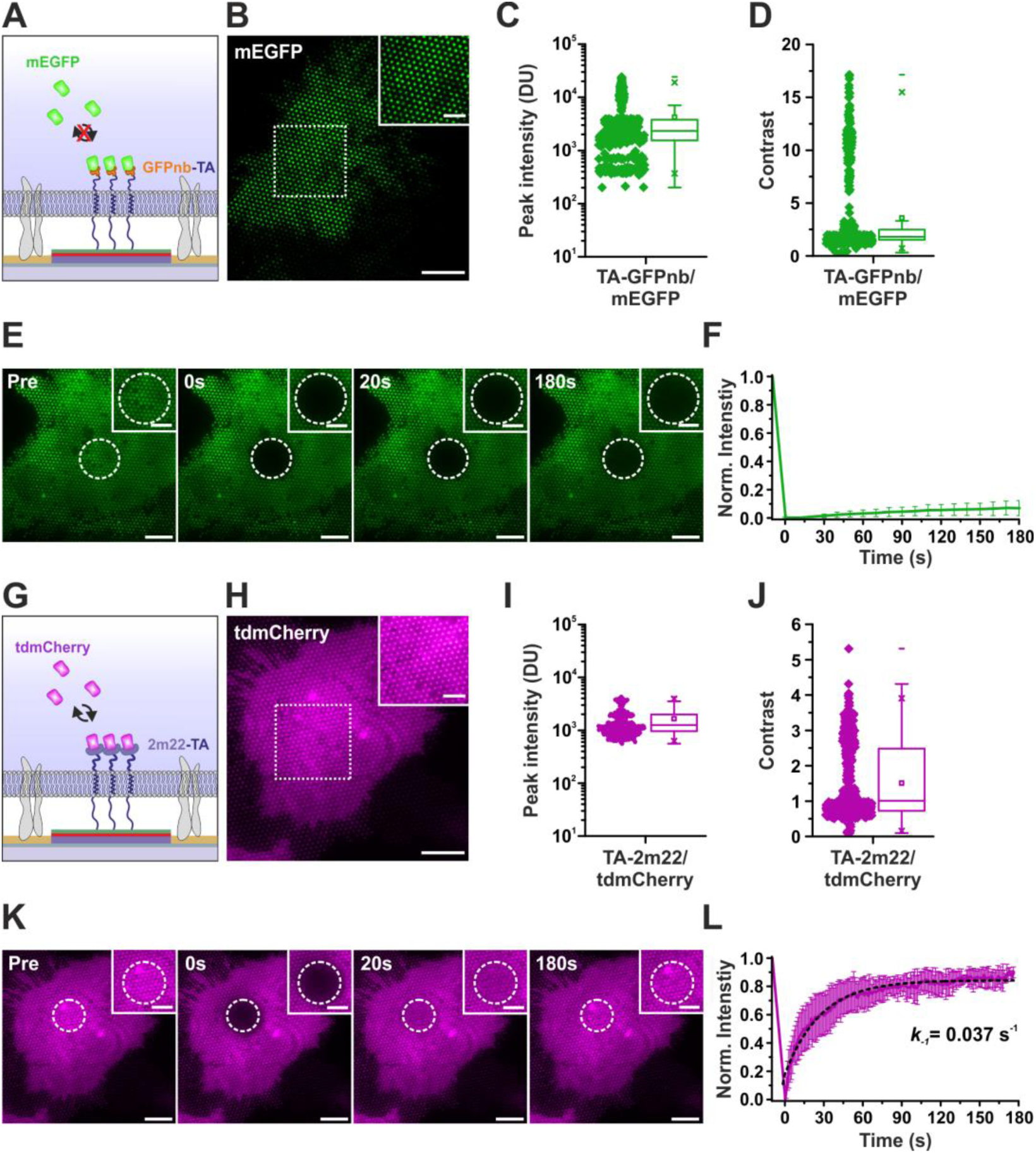
Transmembrane adaptors for cytosolic NDAs in live cells. (A) Schematic representation of the irreversible recruitment of mEGFP through a bifunctional transmembrane adaptor (GFPnb-TA). (B) Representative TIRF microscopy image showing spatially restricted accumulation of cytosolic mEGFP within cNDAs. The dashed box highlights a selected region, and the inset provides a magnified view of individual nanodots. Scale bars: 10 µm; inset: 5 µm. (C, D) Quantitative analysis of peak fluorescence intensity (C) and contrast (D) for GFPnb-TA/mEGFP interactions, verifying efficient immobilization. (E) Time-lapse TIRF microscopy images showing FRAP of mEGFP. Dashed circles indicate the bleached region. Scale bars: 10 µm; insets: 5 µm. (F) Normalized fluorescence intensity recovery curve corresponding to (E). (G) Schematic illustration depicting reversible recruitment of tdmCherry via a bifunctional transmembrane adaptor (2m22-TA). (H) Representative TIRF microscopy image showing the selective capture of tdmCherry within cNDAs. Dashed box marks a selected region, with the inset providing an enlarged visualization of individual nanodots. Scale bars: 10 µm; inset: 5 µm. (I, J) Quantitative analysis of peak fluorescence intensity (I) and contrast (J) for 2m22-TA/tdmCherry. (K) Time-lapse TIRF microscopy images capturing FRAP of tdmCherry. Dashed circles mark the bleached region. Scale bars: 10 µm; insets: 5 µm. (L) Normalized fluorescence intensity recovery curve corresponding to (K). The recovery curve was fitted with a monoexponential function (black dashed line) to quantify the dissociation rate constant (*k_-1_*).

Compared to *in vitro* assays, peak intensity and contrast values of individual nanodots showed substantially broader distribution (**Figure 5**C, D), indicating heterogeneous capturing efficiencies, potentially caused by local differences in membrane-substrate distance. Fluorescence recovery after photobleaching (FRAP) of mEGFP captured into cNDAs showed negligible recovery, in line with a slow exchange kinetics due to the stability of GFPnb/mEGFP complexes (**Figure 5**E, F).^45^ Similarly, efficient capturing of cytosolic tdmCherry into cNDAs was achieved using a corresponding transmembrane adaptor DARPin 2m22 fused to a TMD and an extra-cellular ALFAnb (2m22-TA, **Figure 5**G, H). Peak intensity values of individual nanodots were somewhat lower than mEGFP cNDAs but the contrast values showed large reduction (**Figure 5**I, J), likely to be attributed to higher cytosolic background fluorescence or the lower binding affinity of 2m22:tdmCherry. We therefore anticipated lower complex stability for 2m22:tdmCherry in cNDAs, which was confirmed by FRAP experiments (**Figure 5**K, L). An exchange rate constant of *k_ex_* ≈ 0.04 s^-1^ was obtained by fitting an exponential function, which can be interpreted as the dissociation rate constant *k_-1_* of the 2m22:tdmCherry complex. Given the previously reported *in vitro* affinity of ∼10 nM,^49^ a high on-rate for this complex can be assumed (>10^6^ M^-1^s^-1^).

Leveraging these orthogonal intracellular capturing tools, binary patterning of two distinct cytosolic targets was achieved (Figure S7). For this purpose, PPA and PPS were sequentially CNS-printed onto the same substrate and backfilled with PP-RGD. HeLa cells co-expressing orthogonal transmembrane adaptors 2m22-TA and GFPnb-TMD-SpyCatcher (GFPnb-TS), along with mEGFP and tdmCherry, yielded spatially shifted cNDAs with strong anticorrelation (Figure S7B-F). FRAP experiments confirmed differential exchange dynamics of GFPnb:mEGFP and 2m22:tdmCherry in the binary format (Figure S7G-J). Taken together, these results confirm successful implementation of cNDAs for live cells using two orthogonal transmembrane adaptors with complementary features: fast, high-affinity capturing of GFP-tagged cytosolic bait proteins, and reversible yet rapid capturing of tdmCherry-tagged bait proteins. We here particular focused on leveraging transient intracellular protein capturing by the 2m22 system for probing multiprotein complex formation, which we expect to reduce the exchange dynamics through multivalent interactions thus enhance the selectivity of capturing multiprotein complexes in the cytosol.

### Detection of multimeric protein assemblies in live cell cNDAs

For assessing the capabilities of cNDAs for untangling cooperative interactions in multiprotein complex assembly, we turned to the myddosome complex as a hallmark of supramolecular organizing centers (SMOCs).^50^ Myddosomes have fundamental functions in transmembrane signaling in innate immunity and inflammation in the toll-like receptor/interleukin-1 receptor superfamily.^33, 51,52^ Key player in myddosome assembly is the scaffold protein MyD88, which orchestrates the recruitment and oligomerization of different members of the IL1R-associated kinase (IRAK) family into the signaling complex through death domain (DD)-mediated interactions.^34, 35, 53, 54^ MyD88 together with IRAK4 and IRAK1/IRAK2 supposedly assembles into oligomeric, helical complexes,^34^ which in turn trigger activation of downstream signaling cascades through the ubiquitin ligase TRAF6.^55^ However, previous studies suggested cooperative interactions to be critical for efficient signal propagation,^56^ rendering cNDAs particularly suitable for investigating these processes due to their capability to locally concentrate proteins at high densities within live cells. Given the inherently low endogenous expression levels of myddosome-associated proteins in HeLa cells, we utilized this cell line to ensure precise control over protein assembly. For proof-of-concept experiments, we first explored oligomerization of MyD88 by capturing MyD88 fused to tdmCherry (MyD88-tdmCherry) via the transmembrane adaptor 2m22-TA into cNDAs (**Figure 6**A). Co-expression of MyD88 fused to mEGFP (MyD88-mEGFP) led to strong co-recruitment of both MyD88 fusion proteins into cNDAs, as shown by live-cell TIRF imaging (**Figure 6**B). In contrast, control experiments employing tdmCherry alone did not yield specific co-recruitment, with MyD88-mEGFP instead showing diffuse cytosolic aggregation (**Figure 6**C, D). Quantitative analyses corroborated these observations:

**Figure 6.**
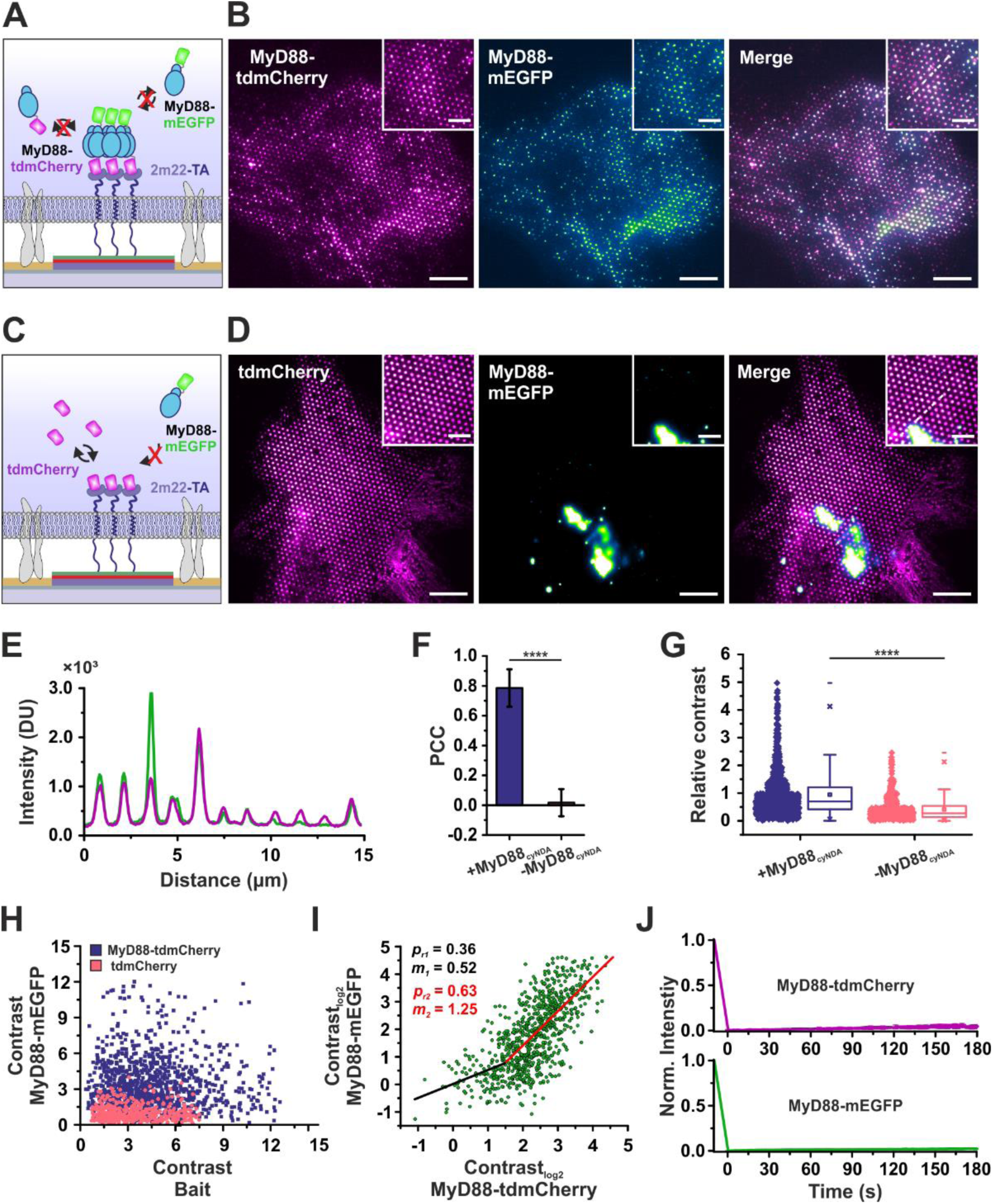
Density-dependent assembly of oligomeric MyD88 complexes in cNDAs. (A) Schematic illustration of MyD88 complex recruitment within cNDAs. MyD88-tdmCherry is captured as bait protein via 2m22-TA, facilitating co-recruitment of MyD88-mEGFP by oligomerization. (B) Representative dual-color TIRF microscopy images of MyD88-tdmCherry (left), corecruited MyD88-mEGFP (middle) and the merged image (right) within cNDAs. Insets provide magnified views of nanodots. White dashed lines indicate regions selected for intensity profiles. Scale bars: 10 µm; insets: 5 µm. (C) Schematic representation of the control experiment where tdmCherry is immobilized. (D) Representative dual-color TIRF microscopy images of the control experiment. Insets provide magnified views of nanodots. White dashed lines indicate regions selected for intensity profiles shown in Figure S8. Scale bars: 10 µm; insets: 5 µm. (E) Fluorescence intensity line profiles for MyD88-tdmCherry (magenta) and MyD88-mEGFP (green) showing overlapping peaks. (F) PCC analysis quantifying the degree of co-localization between MyD88-tdmCherry and MyD88-mEGFP in the presence (+) or absence (-) of immobilized bait (****p < 0.0001). (G) Quantitative comparison of relative contrast values in the presence (+) versus absence (-) of immobilized MyD88-tdmCherry (****p < 0.0001). (H) Scatter plot showing contrast values for individual nanodots from multiple cells. (I) Single-cell correlation analysis between the contrasts of MyD88-tdmCherry and MyD88-mEGFP for approximately 1000 individual nanodots. Pearson’s r-values (p) and slopes (m) are indicated. (J) FRAP analysis showing normalized fluorescence recovery curves for MyD88-tdmCherry (top) and MyD88-mEGFP (bottom).

Pearson’s correlation coefficient (PCC) measurements confirmed significant and specific colocalization of MyD88-tdmCherry and MyD88-mEGFP in cNDAs, whereas this was not the case in control conditions (**Figure 6**E, F, Figure S8A, B). Additionally, comparative analyses of peak intensities and contrasts showed the same result (**Figure 6**G, H, Figure S8C). Detailed analyses at the single-nanodot level demonstrated a biphasic increase of MyD88-mEGFP contrast (prey) as a function of MyD88-tdmCherry density (bait), with Pearson’s correlation coefficient between prey and bait being *p_1_* = 0.36 and *p_2_* = 0.63, respectively (**Figure 6**I). The switch point indicated a critical threshold of MyD88-tdmCherry contrast (2.8) was required to initiate stable MyD88 oligomer formation.

FRAP experiments further confirmed the formation of stable MyD88 oligomers within cNDAs, as no significant fluorescence recovery was detectable for either MyD88-tdmCherry or MyD88-mEGFP (**Figure 6**J and Figure S8F). In contrast, control experiments utilizing tdmCherry alone exhibited the previously shown characteristics of reversible binding (Figure S8D, E). Collectively, these results highlight that cNDAs can steer density-dependent oligomerization of MyD88 into spatially defined and stable supramolecular complexes within live cells.

### Assembly of multiprotein myddosomes in cNDAs

Leveraging spatial control of MyD88 oligomerization in cNDAs, we further explored steps of myddosome assembly. Co-expression of MyD88-tdmCherry and IRAK4 fused to mEGFP (IRAK4-mEGFP) with 2m22-TA confirmed efficient co-recruitment of IRAK4-mEGFP (Figure S9A-D). Correlating the contrast values of MyD88 and IRAK4 at single nanodot level revealed a linear relationship suggesting that seeding of MyD88 oligomers leads to sufficient interaction (Figure S9E, F). Nevertheless, stable oligomerization between MyD88 and IRAK4 within cNDAs was demonstrated by FRAP experiments, confirming their persistent interactions (Figure S9G-J). We next strived for assembly of full myddosome complexes in cNDAs by co-expressing IRAK4-mTagBFP, IRAK1-mEGFP, and TRAF6-tdiRFP together with MyD88-tdmCherry and 2m22-TA (**Figure 7**A). Strikingly, co-recruitment of all four core myddosome components could be unambiguously discerned, with high co-localization and similar peak intensity distributions (**Figure 7**B-E). Correlation analysis of peak intensities at single nanodot level uncovered a more complex picture of myddosome assembly (**Figure 7**F). While the interaction between MyD88 and IRAK4 showed a linear, non-cooperative correlation, we observed pronounced cooperativity between IRAK4 and IRAK1, as well as between MyD88 and IRAK1. Interestingly, the recruitment of TRAF6 was most robust and appeared largely independent of the densities of the remaining components. These analyses provide direct evidence for hierarchical myddosome assembly in a cellular context and demonstrate that individual components engage with distinctly varying degrees of cooperativity.

**Figure 7.**
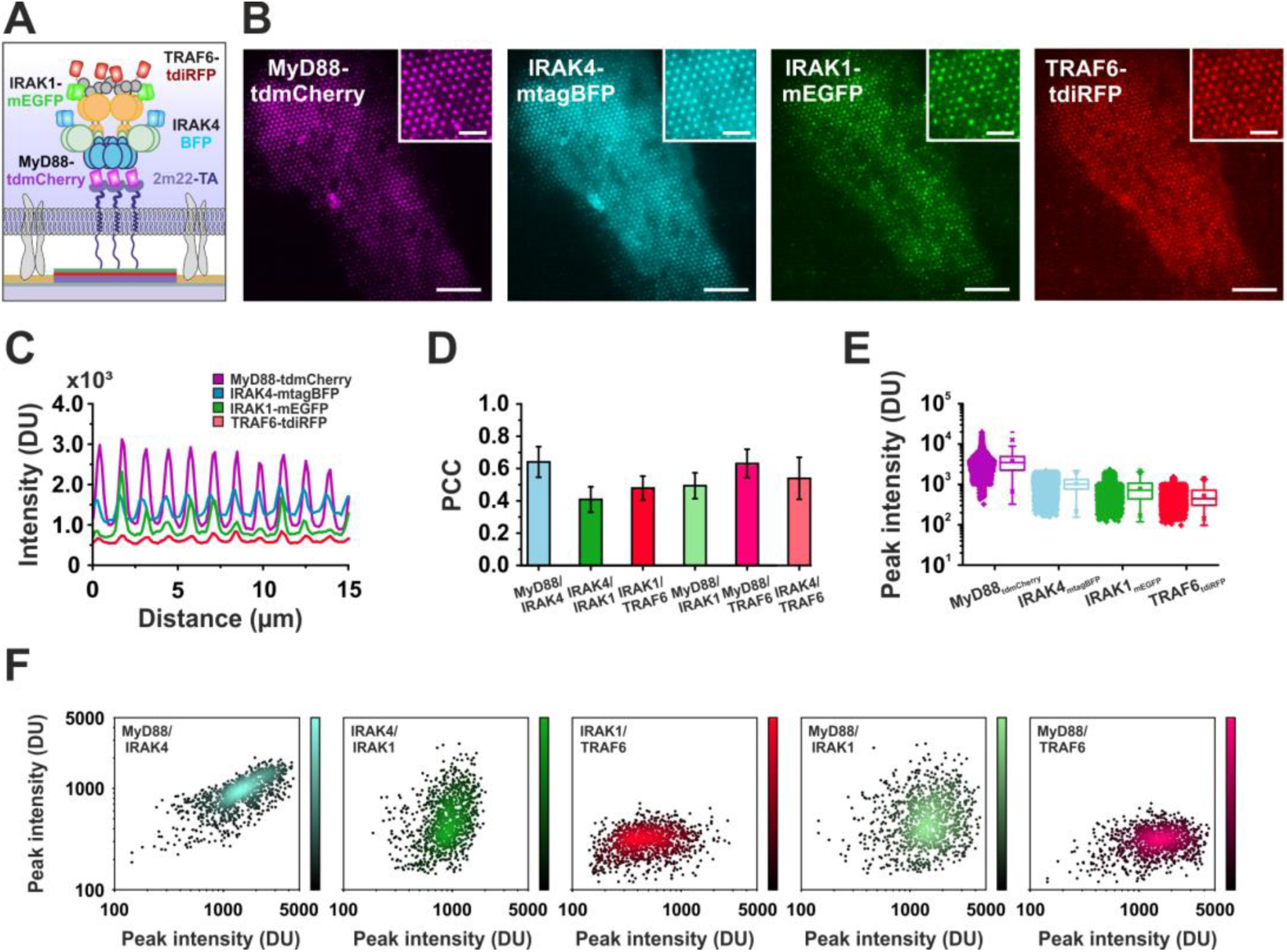
Myddosome formation within cNDAs through co-recruitment of IRAK4, IRAK1, and TRAF6. (A) Schematic illustration depicting the assembly of the myddosome multiprotein complex within cNDAs. Immobilized MyD88-tdmCherry functions as the central scaffold facilitating the hierarchical recruitment of IRAK4-mtagBFP, IRAK1-mEGFP, and TRAF6-tdiRFP. (B) Representative TIRF microscopy images of immobilized MyD88-tdmCherry (magenta) and the co-recruited downstream effectors IRAK4-mTagBFP (cyan), IRAK1-mEGFP (green), and TRAF6-tdiRFP (red). Insets show magnified views of individual nanodots. Scale bars: 10 µm; insets: 5 µm. (C) Fluorescence intensity line profiles of MyD88-tdmCherry, IRAK4-mtagBFP, IRAK1-mEGFP, and TRAF6-tdiRFP within nanodots. (D) PCC quantifying the degree of colocalization between MyD88 and downstream myddosome components (IRAK4, IRAK1, TRAF6). (E) Quantitative comparison of nanodot peak fluorescence intensities across MyD88-tdmCherry, IRAK4-mTagBFP, IRAK1-mEGFP, and TRAF6-tdiRFP. (F) Single-cell correlation analysis of nanodot peak fluorescence intensities between MyD88-IRAK4, IRAK4-IRAK1, IRAK1-TRAF6, MyD88-IRAK1, and MyD88-TRAF6 pairs.

### DNA-PAINT resolves individual myddosome complexes in cNDAs

Having demonstrated that cNDAs can spatially control the assembly of entire myddosomes in live cells, we strived to resolve MyD88 oligomers and their primary interaction partner IRAK4 by DNA-PAINT super-resolution imaging.^36^ To this end, cNDAs containing MyD88-tdmCherry with co-recruited MyD88-mEGFP or IRAK4-mEGFP were stained with commercially available GFPnb conjugated with a docking strand. Representative DNA-PAINT super-resolution images rendered from 40000 frames and the corresponding diffraction-limited images are shown in **Figure 8**A, B and Figure S10 A-D.

**Figure 8.**
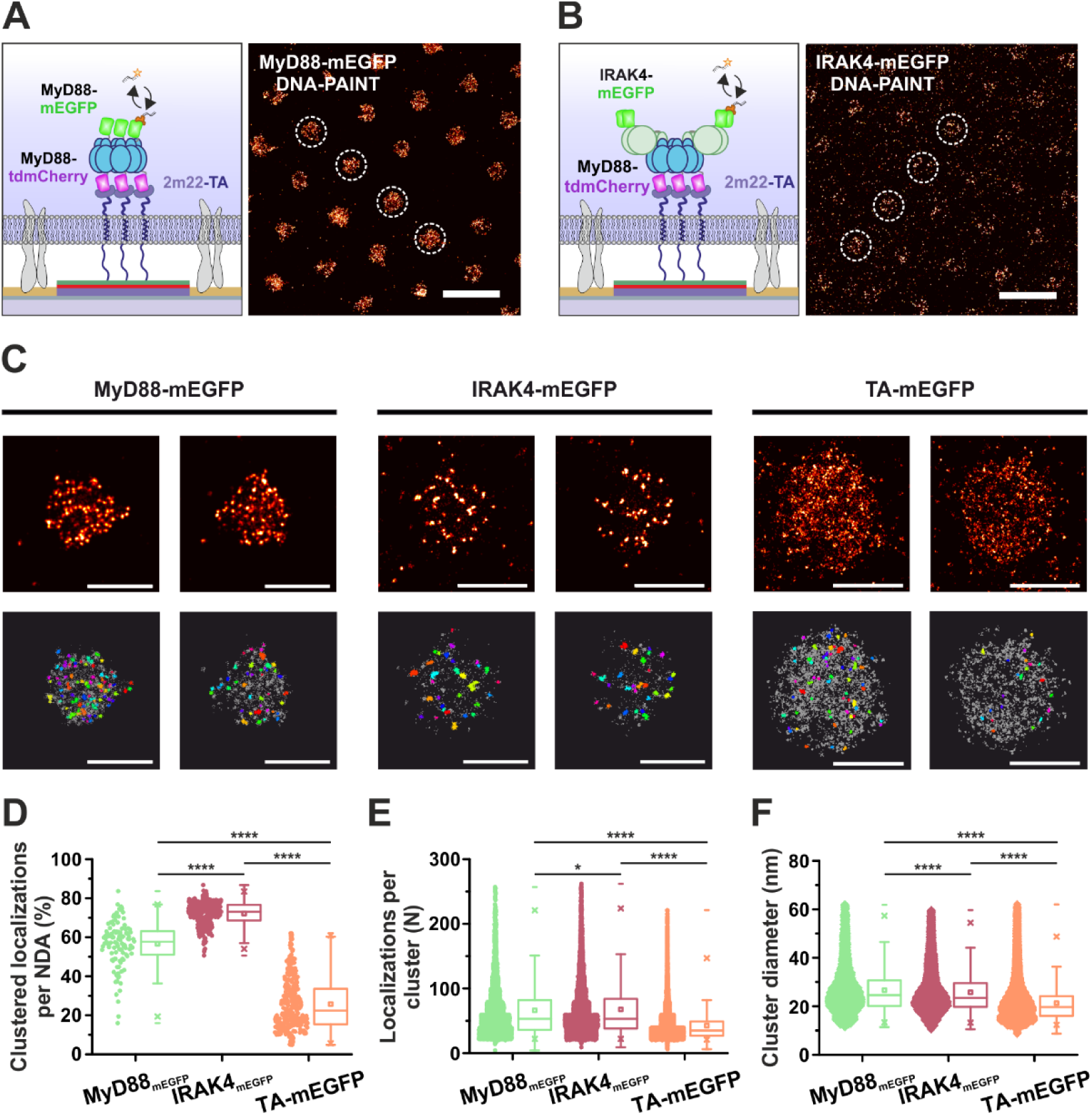
Nanoscale characterization of MyD88-mEGFP and IRAK4-mEGFP clustering within cNDAs using DNA-PAINT super-resolution microscopy. (A) Schematic illustration (left) and representative DNA-PAINT super-resolution image (right) illustrating MyD88-mEGFP clustering within cNDAs. Super-resolution mapping was achieved using an anti-GFP nanobody. Dashed circles highlight nanodots demonstrating distinct clustering patterns. Scale bar: 1 µm. (B) Schematic representation (left) and representative DNA-PAINT super-resolution image (right) illustrating IRAK4-mEGFP clustering within cNDAs. Dashed circles highlight cNDAs demonstrating distinct clustering patterns. Scale bar: 1 µm. (C) DNA-PAINT super-resolution images showing representative clustering characteristics of MyD88-mEGFP (left), IRAK4-mEGFP (middle) and TA-mEGFP (right) within individual nanodots. Upper panels display localization maps, while lower panels depict corresponding DBSCAN-identified clusters. Scale bars: 500 nm. (D-F) Quantitative clustering analysis for MyD88-mEGFP (green), IRAK4-mEGFP (red), and TA-mEGFP (orange). Parameters analyzed include the percentage of clustered localizations per cNDA (D), number of localizations per cluster (E), and cluster diameters (F). Statistical significance: ****p < 0.0001.

Strikingly, strong inhomogeneity and clustering of localizations was observed for both, MyD88 and IRAK4, within cNDAs, but not in control experiments using mEGFP captured via a corresponding transmembrane adaptor (TA-mEGFP, Figure S10E, F) supporting that cNDAs specifically facilitate stable oligomerization of these myddosome components. For both, MyD88 and IRAK4, distinct clusters were identified by DBSCAN cluster analysis, with the majority of localized molecules located within clusters, whereas a much lower level of clustering was detected for the TA-mEGFP control (**Figure 8**C, D). Interestingly, although IRAK4 cNDAs contained overall less localizations per nanodot, the fraction of IRAK4 localizations participating in clusters was higher (∼75 %) as compared to MyD88. This observation is in line with a hierarchical assembly mechanism where scaffold pre-clustering of MyD88 enables IRAK4 co-clustering. One should keep in mind, however, that MyD88 clusters contain both MyD88-tdmCherry and MyD88-mEGFP and only the latter is detected by DNA-PAINT. However, a very similar number of localizations per cluster of ∼ 42 and mean cluster diameters of ∼ 22 nm were found for MyD88 and IRAK4, suggesting correlation of MyD88 and IRAK4 clusters and both substantially exceeding the corresponding data obtained for the mEGFP control (**Figure 8**E, F, Figure S10G, H). These results corroborate that cNDAs effectively steer the hierarchical formation of myddosomes within live cells, thereby opening striking possibilities for imaging individual multiprotein complexes at nanoscale resolution within a cellular context.

### Molecular architecture of myddosomes in cNDAs resolved by RESI

To further push the spatial resolution of DNA-PAINT imaging of myddosomes, we turned to resolution enhancement by sequential imaging (RESI),^57^ focusing on uncovering the molecular organization of IRAK4-mEGFP bound to MyD88-tdmCherry containing cNDAs. After fixation and permeabilization, HeLa cells were stained with an equimolar mixture of anti-GFP nanobodies conjugated with four different, orthogonal docking strands (**Figure 9**A).^57^ Sequential DNA-PAINT with the corresponding imager strands yielded four independent localization datasets. Representative SR images are shown in **Figure 9**B in different rendering formats: when all four localization data sets were rendered together, a similar SR image similar to classic DNA-PAINT of IRAK4-mEGFP was observed (**Figure 9**B, left image); color-coding individual localization data sets clearly identified spatially separated localization cluster corresponding to individual IRAK4 molecules (**Figure 9**B, center image).

**Figure 9.**
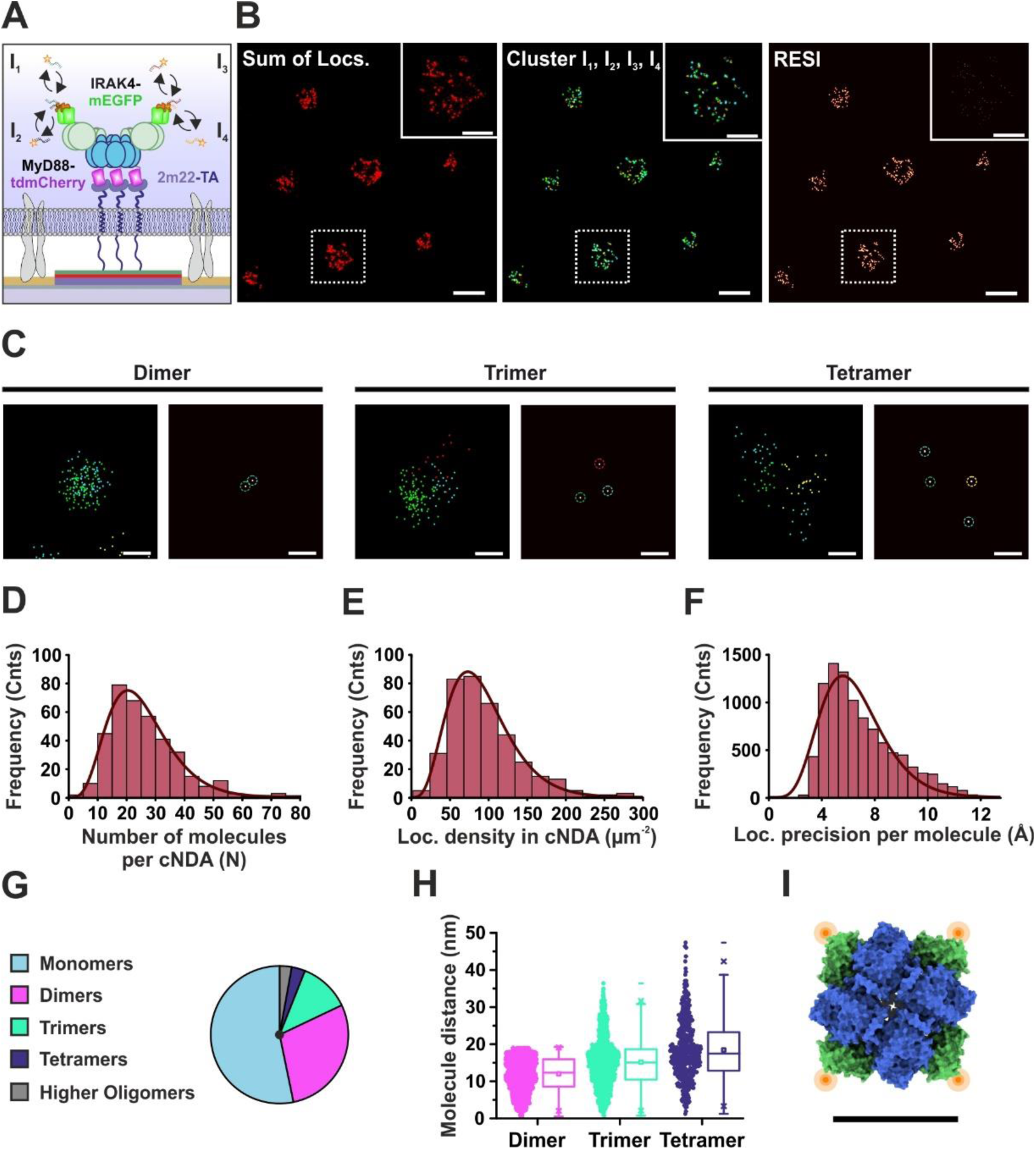
RESI of IRAK4-mEGFP resolves spatial organization at sub-nanometer precision. (A) Schematic illustration of the RESI approach applied to IRAK4-mEGFP within cNDAs. Sequential imaging rounds (I₁ to I₄) capture distinct localization events, enabling enhanced spatial resolution to resolve discrete molecular assemblies. (B) Representative RESI reconstruction showing cumulative localizations (left), sequentially resolved individual clusters from separate imaging rounds (middle), and the final RESI-processed image (right). Insets provide detailed views of resolved oligomeric clusters. Scale bars: 500 nm; insets: 200 nm. (C) Classification of IRAK4-mEGFP oligomers within nanodots using RESI. Examples illustrate identified dimers, trimers, and tetramers, alongside corresponding localization maps. Scale bars: 10 nm. (D-F) Quantitative RESI analysis of IRAK4-mEGFP molecules per cNDA showing: (D) the distribution of molecule numbers per nanodot, (E) localization density per nanodot, and (F) localization precision per molecule. All histograms were fitted using Gamma distributions. (G) Population analysis of IRAK4-mEGFP depicting the fractions of monomers, dimers, trimers, tetramers, and higher-order oligomers. (H) Intermolecular distance analysis within different IRAK4-mEGFP oligomers. (I) AlphaFold3 structural model of 4× IRAK4-mEGFP (blue: IRAK4, green: mEGFP) together with death domains of MyD88 (6×, not shown here) and IRAK1 (4×, not shown here). Approximate localization of the fluorophores used for RESI imaging is depicted in orange. Scale bar: 10 nm.

Rendering of these clusters yielded a reconstructed RESI image representing the molecular arrangement of IRAK4 within myddosomes (**Figure 9**B, right image). Further analysis of these RESI images revealed heterogeneous, but well-defined oligomeric assemblies of IRAK4, including discrete dimers, trimers, and less frequently, tetramers (**Figure 9**C). Quantitative analysis identified a mean number of ∼ 20 IRAK4 molecules per cNDA with a mean molecular density of ∼ 73 molecules/µm² (**Figure 9**D, E). The exceptionally high localization precision achieved by RESI (< 1 nm) (**Figure 9**F) confirmed the potential of this method to resolve the structural organization of myddosomes, models of which show a typical diameter of > 10 nm (Figure S11). Oligomer analysis further revealed a high level of dimers and trimers with mean distances of 12-15 nm, as well as a small fraction of tetramers, which only showed slightly increased intermolecular distance (**Figure 9**G, H). These findings are in striking agreement with structural models of IRAK4 tetramers in a myddosome as predicted by AlphaFold3 (**Figure 9**H).^58^ Given the theoretical maximum labeling efficiency of ∼ 60 % and the stochastic labeling with the four different docking strands, the probability of detecting ground-truth tetramers is rather low, while dimers and trimers are expected to be dominating. Overall, these proof-of-concept experiments highlight that RESI imaging in combination with cNDAs enables resolving individual protein complexes with molecular precision and provides direct experimental access to the structural organization of supramolecular assemblies in the native cellular context.

## Conclusions

The diversity and plasticity of protein function in cells relates to their ability to dynamically assemble into complexes with different composition and stoichiometry^59–61^ which is dictated by the physiological context. However, systematic analyses of protein complexes in cells remain highly challenging, as generic tools for controlling their assembly in the physiological context are lacking. Here, we introduce a comprehensive toolbox for efficient spatial control of cytosolic protein complex assemblies by live-cell cytosolic nanodot arrays (cNDAs). Using synthetic polymer carriers with rapid binding functionalization for capillary nanostamping onto hydrophobic substrates, protein capturing efficiency could be enhanced by more than an order of magnitude, while maintaining nanodot size substantially below 500 nm. Characterization of the bNDA nanoscale architecture by DNA-PAINT highlighted the improved functionalization density and ultrahigh contrast were achieved under optimized conditions. Introducing orthogonal surface functionalization groups further enhanced the versatility, enabling controlled co-organization of two distinct proteins at nanoscopic and at molecular scale, which was demonstrated to be compatible with the application for probing cytosolic protein-protein interactions in live cells. Here, we focused on assessing the capabilities of cNDAs for studying the assembly of multi-protein complexes, employing the myddosome as a model system. We demonstrate that high local densities of bait proteins (MyD88) achieved by optimizing adaptor’s affinity in cNDAs enabled to locally initiate protein oligomerization, while allowing additional homo- and heteromeric interaction partners (IRAK4, IRAK1 and TRAF6) to be efficiently recruited. The ability to correlate the density of different interaction partners at single nanodot level provided a detailed picture of the cooperativity involved in hierarchical multiprotein assemblies. Finally, by combining DNA-PAINT super-resolution imaging techniques with cNDAs, we demonstrated ultrastructural analysis of multiprotein complexes with resolution down to the sub-nanometer level. Detailed cluster analysis from classic and RESI imaging mode confirmed the assembly of relatively well-defined myddosomes in the cellular context, with stoichiometry and architecture aligning with current concepts and AlphaFold3 models. We here limited our study to proof-of-concept experiments with structural analysis of only partially assembled myddosomes. Follow-up studies using multiplexed DNA-PAINT, quantitative PAINT and RESI imaging, exploiting new tools continually developed in the field, will enable ultrastructural characterization of fully assembled myddosomes. Our initial analyses suggest that myddosome assembly in cNDAs is oriented perpendicular to the plasma membrane. Thus, three-dimensional imaging methods with high axial resolution (e.g. MIET-PAINT)^62^ can further unveil the architecture of these complexes in live cells. Moreover, cNDAs can be integrated with optogenetic actuators or microfluidic modules which will allow precise control of stimuli for time-resolved dissection of signaling cascades inside cells.

## Data and materials availability

All data needed to evaluate the conclusions in the paper are present in the paper and/or the Supplementary Materials. Additional data related to this paper may be requested from the authors.

## Acknowledgement

We thank Marcus Taylor at Max Planck Institute for Infection Biology, Berlin for providing the IRAK1 and IRAK4 plasmids. We thank A. Budke-Gieseking, G. Hikade, H. Kenneweg and W. Kohl for technical support. This work was supported by the Deutsche Forschungsgemeinschaft through grants SFB 1557 (467522186) to J.P. and R.K., RTG 2900 (501879556) to J.P., M.S., and C.Y., PI 405/9-2 (272553338) and PI 405/15-2 (326558201) to J.P., and YO 166/1-1 (405319862) to C.Y. J.P. acknowledges intramural funding from the Lower Saxony Ministry of Science and Culture.

## Author Contributions

J.P. and C.Y. conceived the project together with A.F.. A.F. prepared the bNDAs and cNDAs, performed the experiments of myddosome and conducted data analysis. M.P. contributed to synthesis of the carrier polymers and the functional assays on surface. M.H. contributed to DNA-PAINT imaging. C.D. contributed to surface silanization. E.S. conducted the characterization of silanized surfaces under the supervision of M.S.. R.K. contributed to super-resolution imaging and image evaluations. A.F., C.Y. and J.P. wrote the manuscript with contributions from all authors.

## References

(1) Nooren, I. M. A.; Thornton, J. M. Diversity of protein–protein interactions. The EMBO Journal 2003, 22 (14), 3486–3492. DOI: 10.1093/emboj/cdg359.

(2) Jones, S.; Thornton, J. M. Principles of protein-protein interactions. Proceedings of the National Academy of Sciences 1996, 93 (1), 13–20. DOI: doi:10.1073/pnas.93.1.13.

(3) Perkins, J. R.; Diboun, I.; Dessailly, B. H.; Lees, J. G.; Orengo, C. Transient Protein-Protein Interactions: Structural, Functional, and Network Properties. Structure 2010, 18 (10), 1233–1243. DOI: 10.1016/j.str.2010.08.007.

(4) Kapoor, A.; Mondal, S.; Chaudhary, A.; Sharma, S.; Mehra, P.; Prasad, A. A topological review on protein–protein interactions: the development and promises in the era of omics. Journal of Proteins and Proteomics 2024, 15 (3), 523–544. DOI: 10.1007/s42485-024-00160-w.

(5) Michaelis, A. C.; Brunner, A. D.; Zwiebel, M.; Meier, F.; Strauss, M. T.; Bludau, I.; Mann, M. The social and structural architecture of the yeast protein interactome. Nature 2023, 624 (7990), 192–200. DOI: 10.1038/s41586-023-06739-5.

(6) Liu, X.; Abad, L.; Chatterjee, L.; Cristea, I. M.; Varjosalo, M. Mapping protein-protein interactions by mass spectrometry. Mass Spectrom Rev 2024. DOI: 10.1002/mas.21887.

(7) Rao, V. S.; Srinivas, K.; Sujini, G. N.; Kumar, G. N. Protein-protein interaction detection: methods and analysis. Int J Proteomics 2014, 2014, 147648. DOI: 10.1155/2014/147648 .

(8) Mangiarotti, A.; Siri, M.; Tam, N. W.; Zhao, Z.; Malacrida, L.; Dimova, R. Biomolecular condensates modulate membrane lipid packing and hydration. Nature Communications 2023, 14 (1), 6081. DOI: 10.1038/s41467-023-41709-5.

(9) Kim, N.; Yun, H.; Lee, H.; Yoo, J.-Y. Interplay between membranes and biomolecular condensates in the regulation of membrane-associated cellular processes. Experimental & Molecular Medicine 2024, 56 (11), 2357–2364. DOI: 10.1038/s12276-024-01337-5.

(10) Alom Ruiz, S.; Chen, C. S. Microcontact printing: a tool to pattern. Soft Matter 2007, 3, 168.

(11) Li, Y.; Jiang, W.; Zhou, X.; Long, Y.; Sun, Y.; Zeng, Y.; Yao, X. Advances in Regulating Cellular Behavior Using Micropatterns. Yale J Biol Med 2023, 96 (4), 527–547. DOI: 10.59249/uxoh1740.

(12) Karimian, T.; Lanzerstorfer, P.; Weghuber, J. Soft lithography-based biomolecule patterning techniques and their applications in subcellular protein interaction analysis. Materials Today Bio 2025, 32, 101672. DOI: 10.1016/j.mtbio.2025.101672.

(13) Mossman, K. D.; Campi, G.; Groves, J. T.; Dustin, M. L. Altered TCR Signaling from Geometrically Repatterned Immunological Synapses. Science 2005, 310 (5751), 1191–1193. DOI: doi:10.1126/science.1119238.

(14) Schwarzenbacher, M.; Kaltenbrunner, M.; Brameshuber, M.; Hesch, C.; Paster, W.; Weghuber, J.; Heise, B.; Sonnleitner, A.; Stockinger, H.; Schütz, G. J. Micropatterning for quantitative analysis of protein-protein interactions in living cells. Nat Methods 2008, 5 (12), 1053–1060. DOI: 10.1038/nmeth.1268.

(15) Deeg, J.; Axmann, M.; Matic, J.; Liapis, A.; Depoil, D.; Afrose, J.; Curado, S.; Dustin, M. L.; Spatz, J. P. T Cell Activation is Determined by the Number of Presented Antigens. Nano Letters 2013, 13 (11), 5619–5626. DOI: 10.1021/nl403266t.

(16) Lochte, S.; Waichman, S.; Beutel, O.; You, C.; Piehler, J. Live cell micropatterning reveals the dynamics of signaling complexes at the plasma membrane. J Cell Biol 2014, 207 (3), 407–418. DOI: 10.1083/jcb.201406032.

(17) Wedeking, T.; Löchte, S.; Birkholz, O.; Wallenstein, A.; Trahe, J.; Klingauf, J.; Piehler, J.; You, C. Spatiotemporally Controlled Reorganization of Signaling Complexes in the Plasma Membrane of Living Cells. Small 2015, 11 (44), 5912–5918. DOI: 10.1002/smll.201502132.

(18) Sevcsik, E.; Brameshuber, M.; Fölser, M.; Weghuber, J.; Honigmann, A.; Schütz, G. J. GPI-anchored proteins do not reside in ordered domains in the live cell plasma membrane. Nature Communications 2015, 6 (1), 6969. DOI: 10.1038/ncomms7969.

(19) Cai, H.; Muller, J.; Depoil, D.; Mayya, V.; Sheetz, M. P.; Dustin, M. L.; Wind, S. J. Full control of ligand positioning reveals spatial thresholds for T cell receptor triggering. Nature Nanotechnology 2018, 13 (7), 610–617. DOI: 10.1038/s41565-018-0113-3.

(20) Lu, C.-H.; Pedram, K.; Tsai, C.-T.; Jones, T.; Li, X.; Nakamoto, M. L.; Bertozzi, C. R.; Cui, B. Membrane curvature regulates the spatial distribution of bulky glycoproteins. Nature Communications 2022, 13 (1), 3093. DOI: 10.1038/s41467-022-30610-2.

(21) Mayer, I.; Karimian, T.; Gordiyenko, K.; Angelin, A.; Kumar, R.; Hirtz, M.; Mikut, R.; Reischl, M.; Stegmaier, J.; Zhou, L.;, et al. Surface-Patterned DNA Origami Rulers Reveal Nanoscale Distance Dependency of the Epidermal Growth Factor Receptor Activation. Nano Letters 2024, 24 (5), 1611–1619. DOI: 10.1021/acs.nanolett.3c04272.

(22) Gandor, S.; Reisewitz, S.; Venkatachalapathy, M.; Arrabito, G.; Reibner, M.; Schroder, H.; Ruf, K.; Niemeyer, C. M.; Bastiaens, P. I.; Dehmelt, L. A protein-interaction array inside a living cell. Angew Chem Int Ed Engl 2013, 52 (18), 4790–4794. DOI: 10.1002/anie.201209127.

(23) Bag, N.; Huang, S.; Wohland, T. Plasma membrane organization of epidermal growth factor receptor in resting and ligand-bound states. Biophys. J. 2015, 109, 1925.

(24) Arimoto, K. I.; Löchte, S.; Stoner, S. A.; Burkart, C.; Zhang, Y.; Miyauchi, S.; Wilmes, S.; Fan, J. B.; Heinisch, J. J.; Li, Z.;, et al. STAT2 is an essential adaptor in USP18-mediated suppression of type I interferon signaling. Nat Struct Mol Biol 2017, 24 (3), 279–289. DOI: 10.1038/nsmb.3378.

(25) Chen, Z.; Oh, D.; Biswas, K. H.; Yu, C.-H.; Zaidel-Bar, R.; Groves, J. T. Spatially modulated ephrinA1:EphA2 signaling increases local contractility and global focal adhesion dynamics to promote cell motility. Proceedings of the National Academy of Sciences 2018, 115 (25), E5696–E5705. DOI: 10.1073/pnas.1719961115.

(26) Huang, W. Y. C.; Alvarez, S.; Kondo, Y.; Lee, Y. K.; Chung, J. K.; Lam, H. Y. M.; Biswas, K. H.; Kuriyan, J.; Groves, J. T. A molecular assembly phase transition and kinetic proofreading modulate Ras activation by SOS. Science 2019, 363 (6431), 1098–1103. DOI: 10.1126/science.aau5721.

(27) Chen, Z.; Oh, D.; Biswas, K. H.; Zaidel-Bar, R.; Groves, J. T. Probing the effect of clustering on EphA2 receptor signaling efficiency by subcellular control of ligand-receptor mobility. eLife 2021, 10, e67379. DOI: 10.7554/eLife.67379.

(28) Hager, R.; Müller, U.; Ollinger, N.; Weghuber, J.; Lanzerstorfer, P. Subcellular Dynamic Immunopatterning of Cytosolic Protein Complexes on Microstructured Polymer Substrates. ACS Sensors 2021, 6 (11), 4076–4088. DOI: 10.1021/acssensors.1c01574.

(29) Sánchez, M. F.; Faria, S.; Frühschulz, S.; Werkmann, L.; Winter, C.; Karimian, T.; Lanzerstorfer, P.; Plochberger, B.; Weghuber, J.; Tampé, R. Engineering Mesoscale T Cell Receptor Clustering by Plug-and-Play Nanotools. Advanced Materials 2024, 36 (45), 2310407. DOI: 10.1002/adma.202310407.

(30) Philippi, M.; You, C.; Richter, C. P.; Schmidt, M.; Thien, J.; Lisse, D.; Wollschlager, J.; Piehler, J.; Steinhart, M. Close-packed silane nanodot arrays by capillary nanostamping coupled with heterocyclic silane ring opening. RSC Adv 2019, 9 (43), 24742–24750. DOI: 10.1039/c9ra03440d.

(31) Philippi, M.; Richter, C. P.; Kappen, M.; Watrinet, I.; Miao, Y.; Runge, M.; Jorde, L.; Korneev, S.; Holtmannspotter, M.; Kurre, R.;, et al. Biofunctional Nanodot Arrays in Living Cells Uncover Synergistic Co-Condensation of Wnt Signalodroplets. Small 2022, 18 (50), e2203723. DOI: 10.1002/smll.202203723 From NLM Medline.

(32) Cao, F.; Deliz-Aguirre, R.; Gerpott, F. H.; Ziska, E.; Taylor, M. J. Myddosome clustering in IL-1 receptor signaling regulates the formation of an NF-kB activating signalosome. EMBO reports 2023, 24 (10), e57233. DOI: 10.15252/embr.202357233.

(33) Fisch, D.; Zhang, T.; Sun, H.; Ma, W.; Tan, Y.; Gygi, S. P.; Higgins, D. E.; Kagan, J. C. Molecular definition of the endogenous Toll-like receptor signalling pathways. Nature 2024, 631 (8021), 635–644. DOI: 10.1038/s41586-024-07614-7.

(34) Lin, S. C.; Lo, Y. C.; Wu, H. Helical assembly in the MyD88-IRAK4-IRAK2 complex in TLR/IL-1R signalling. Nature 2010, 465 (7300), 885–890. DOI: 10.1038/nature09121.

(35) Moncrieffe, M. C.; Bollschweiler, D.; Li, B.; Penczek, P. A.; Hopkins, L.; Bryant, C. E.; Klenerman, D.; Gay, N. J. MyD88 Death-Domain Oligomerization Determines Myddosome Architecture: Implications for Toll-like Receptor Signaling. Structure 2020, 28 (3), 281–289.e283. DOI: 10.1016/j.str.2020.01.003.

(36) Schnitzbauer, J.; Strauss, M. T.; Schlichthaerle, T.; Schueder, F.; Jungmann, R. Super-resolution microscopy with DNA-PAINT. Nature Protocols 2017, 12 (6), 1198–1228. DOI: 10.1038/nprot.2017.024.

(37) VandeVondele, S.; Voros, J.; Hubbell, J. A. RGD-grafted poly-L-lysine-graft-(polyethylene glycol) copolymers block non-specific protein adsorption while promoting cell adhesion. Biotechnol Bioeng 2003, 82 (7), 784–790. DOI: 10.1002/bit.10625.

(38) Ruiz-Taylor, L. A.; Martin, T. L.; Zaugg, F. G.; Witte, K.; Indermuhle, P.; Nock, S.; Wagner, P. Monolayers of derivatized poly(l-lysine)-grafted poly(ethylene glycol) on metal oxides as a class of biomolecular interfaces. Proceedings of the National Academy of Sciences 2001, 98 (3), 852–857. DOI: doi:10.1073/pnas.98.3.852.

(39) Wang, C.; Yang, H.; Chen, F.; Peng, L.; Gao, H.-f.; Zhao, L.-p. Influences of VTMS/SiO2 ratios on the contact angle and morphology of modified super-hydrophobic silicon dioxide material by vinyl trimethoxy silane. Results in Physics 2018, 10, 891–902. DOI: 10.1016/j.rinp.2018.08.007.

(40) Los, G. V.; Encell, L. P.; McDougall, M. G.; Hartzell, D. D.; Karassina, N.; Zimprich, C.; Wood, M. G.; Learish, R.; Ohana, R. F.; Urh, M.;, et al. HaloTag: a novel protein labeling technology for cell imaging and protein analysis. ACS Chem Biol 2008, 3 (6), 373–382. DOI: 10.1021/cb800025k.

(41) Götzke, H.; Kilisch, M.; Martínez-Carranza, M.; Sograte-Idrissi, S.; Rajavel, A.; Schlichthaerle, T.; Engels, N.; Jungmann, R.; Stenmark, P.; Opazo, F.;, et al. The ALFA-tag is a highly versatile tool for nanobody-based bioscience applications. Nature Communications 2019, 10 (1), 4403. DOI: 10.1038/s41467-019-12301-7.

(42) Keeble, A. H.; Turkki, P.; Stokes, S.; Khairil Anuar, I. N. A.; Rahikainen, R.; Hytönen, V. P.; Howarth, M. Approaching infinite affinity through engineering of peptide–protein interaction. Proceedings of the National Academy of Sciences 2019, 116 (52), 26523–26533. DOI: 10.1073/pnas.1909653116.

(43) Lisse, D.; Wilkens, V.; You, C.; Busch, K.; Piehler, J. Selective targeting of fluorescent nanoparticles to proteins inside live cells. Angew Chem Int Ed Engl 2011, 50 (40), 9352–9355. DOI: 10.1002/anie.201101499.

(44) Gavutis, M.; Lata, S.; Lamken, P.; Muller, P.; Piehler, J. Lateral ligand-receptor interactions on membranes probed by simultaneous fluorescence-interference detection. Biophys J 2005, 88 (6), 4289–4302. DOI: 10.1529/biophysj.104.055855.

(45) Rothbauer, U.; Zolghadr, K.; Muyldermans, S.; Schepers, A.; Cardoso, M. C.; Leonhardt, H. A versatile nanotrap for biochemical and functional studies with fluorescent fusion proteins. Mol Cell Proteomics 2008, 7 (2), 282–289. DOI: 10.1074/mcp.M700342-MCP200.

(46) Schmidt, M.; Philippi, M.; Münzner, M.; Stangl, J. M.; Wieczorek, R.; Harneit, W.; Müller-Buschbaum, K.; Enke, D.; Steinhart, M. Capillary Nanostamping with Spongy Mesoporous Silica Stamps. Advanced Functional Materials 2018, 28 (23), 1800700. DOI: 10.1002/adfm.201800700.

(47) Plückthun, A. Designed ankyrin repeat proteins (DARPins): binding proteins for research, diagnostics, and therapy. Annu Rev Pharmacol Toxicol 2015, 55, 489–511. DOI: 10.1146/annurev-pharmtox-010611-134654.

(48) Hansen, S.; Stüber, J. C.; Ernst, P.; Koch, A.; Bojar, D.; Batyuk, A.; Plückthun, A. Design and applications of a clamp for Green Fluorescent Protein with picomolar affinity. In Sci Rep, 2017; Vol. 7, p 16292.

(49) Brauchle, M.; Hansen, S.; Caussinus, E.; Lenard, A.; Ochoa-Espinosa, A.; Scholz, O.; Sprecher, S. G.; Pluckthun, A.; Affolter, M. Protein interference applications in cellular and developmental biology using DARPins that recognize GFP and mCherry. Biology open 2014, 3 (12), 1252–1261. DOI: 10.1242/bio.201410041.

(50) Kagan, J. C.; Magupalli, V. G.; Wu, H. SMOCs: supramolecular organizing centres that control innate immunity. Nat Rev Immunol 2014, 14 (12), 821–826. DOI: 10.1038/nri3757.

(51) Pereira, M.; Gazzinelli, R. T. Regulation of innate immune signaling by IRAK proteins. Frontiers in Immunology 2023, 14, Review. DOI: 10.3389/fimmu.2023.1133354.

(52) Balka, K. R.; De Nardo, D. Understanding early TLR signaling through the Myddosome. Journal of Leukocyte Biology 2019, 105 (2), 339–351. DOI: 10.1002/JLB.MR0318-096R.

(53) Motshwene, P. G.; Moncrieffe, M. C.; Grossmann, J. G.; Kao, C.; Ayaluru, M.; Sandercock, A. M.; Robinson, C. V.; Latz, E.; Gay, N. J. An oligomeric signaling platform formed by the Toll-like receptor signal transducers MyD88 and IRAK-4. J Biol Chem 2009, 284 (37), 25404–25411. DOI: 10.1074/jbc.M109.022392.

(54) Gay, N. J.; Symmons, M. F.; Gangloff, M.; Bryant, C. E. Assembly and localization of Toll-like receptor signalling complexes. Nat Rev Immunol 2014, 14 (8), 546–558. DOI: 10.1038/nri3713.

(55) Ye, H.; Arron, J. R.; Lamothe, B.; Cirilli, M.; Kobayashi, T.; Shevde, N. K.; Segal, D.; Dzivenu, O. K.; Vologodskaia, M.; Yim, M.;, et al. Distinct molecular mechanism for initiating TRAF6 signalling. Nature 2002, 418 (6896), 443–447. DOI: 10.1038/nature00888.

(56) Deliz-Aguirre, R.; Cao, F.; Gerpott, F. H. U.; Auevechanichkul, N.; Chupanova, M.; Mun, Y.; Ziska, E.; Taylor, M. J. MyD88 oligomer size functions as a physical threshold to trigger IL1R Myddosome signaling. The Journal of cell biology 2021, 220 (7). DOI: 10.1083/jcb.202012071.

(57) Reinhardt, S. C. M.; Masullo, L. A.; Baudrexel, I.; Steen, P. R.; Kowalewski, R.; Eklund, A. S.; Strauss, S.; Unterauer, E. M.; Schlichthaerle, T.; Strauss, M. T.;, et al. Ångström-resolution fluorescence microscopy. Nature 2023, 617 (7962), 711–716. DOI: 10.1038/s41586-023-05925-9.

(58) Abramson, J.; Adler, J.; Dunger, J.; Evans, R.; Green, T.; Pritzel, A.; Ronneberger, O.; Willmore, L.; Ballard, A. J.; Bambrick, J.;, et al. Accurate structure prediction of biomolecular interactions with AlphaFold 3. Nature 2024, 630 (8016), 493–500. DOI: 10.1038/s41586-024-07487-w.

(59) Alarcón, B.; Swamy, M.; van Santen, H. M.; Schamel, W. W. A. T-cell antigen-receptor stoichiometry: pre-clustering for sensitivity. EMBO reports 2006, 7 (5), 490–495. DOI: 10.1038/sj.embor.7400682.

(60) Case, L. B.; Zhang, X.; Ditlev, J. A.; Rosen, M. K. Stoichiometry controls activity of phase-separated clusters of actin signaling proteins. Science 2019, 363 (6431), 1093–1097. DOI: 10.1126/science.aau6313.

(61) Tveriakhina, L.; Scanavachi, G.; Egan, E. D.; Da Cunha Correia, R. B.; Martin, A. P.; Rogers, J. M.; Yodh, J. S.; Aster, J. C.; Kirchhausen, T.; Blacklow, S. C. Temporal dynamics and stoichiometry in human Notch signaling from Notch synaptic complex formation to nuclear entry of the Notch intracellular domain. Developmental Cell 2024, 59 (11), 1425–1438.e1428. DOI: 10.1016/j.devcel.2024.03.021.

(62) Oleksiievets, N.; Mougios, N.; Jans, D. C.; Hauke, L.; Thiele, J. C.; Basak, S.; Jakobs, S.; Opazo, F.; Enderlein, J.; Tsukanov, R. Three-dimensional multi-target super-resolution microscopy of cells using Metal-Induced Energy Transfer and DNA-PAINT. *bioRxiv* 2024, 2024.2004.2002.587536. DOI: 10.1101/2024.04.02.587536.

