## Supporting Information for "A Versatile Toolbox for Nanoscale Interrogation of Multiprotein Assemblies inside Living Cells"

Supplementary References ………… page 12

Supplementary Figures S1-S11 ………… pages 13-24

Supplementary Tables S1-S7 ………… pages 25-28

### Materials and Methods

#### Synthesis of functionalized carrier polymers

PLL-PEG conjugates functionalized with ALFAtag (PLL-PEG-ALFAtag) or SpyTag003 (PLL-PEG-SpyTag) were synthesized through a two-step coupling reaction. Briefly, cysteine-containing peptides (Ac-CPSRLEEELRRRLTE for ALFAtag; RGVPHIVMVDAYKRYKAC for SpyTag) (12 µmol, 23.2 mg) were dissolved in 100 µL HBS buffer (100 mM NaCl, 100 mM HEPES, pH 7.5). Subsequently, MAL-PEG$_{3k}$-NHS (12 µmol, 36 mg), pre-dissolved in 100 µL HBS buffer, was added to initiate maleimide-thiol coupling. The reaction proceeded for 15 min at room temperature under gentle shaking. Next, poly-L-lysine hydrobromide (molecular weight: 15 - 30 kDa, 7.5 mg), dissolved in 300 µL HBS buffer, was added to the mixture. The conjugation reaction was carried out overnight at room temperature with continuous stirring. The resulting conjugates were purified by dialysis (14 kDa molecular weight cut-off) against Milli-Q water (2 L, 24 h), followed by lyophilization to yield functionalized PLL-PEG conjugates as white powder. BSA-based conjugates (BSA-HTL and BSA-ALFAtag), as well as PLL-PEG-HTL, were synthesized according to established protocols described previously.[1, 2]

#### Surface silanization of glass substrates

Glass substrates and silica transducers used for Reflectance Interference Spectroscopy (RIFS) were initially cleaned by sequential rinsing with isopropanol and Milli-Q water to remove surface contaminants. Subsequently, substrates underwent surface activation by oxygen plasma treatment for 10 min using a Diener Femto plasma cleaner. Following activation, surfaces were immediately silanized: For silanization with (4-chlorophenyl)triethoxysilane (CPTES), activated substrates were immersed into a 2 % (v/v) CPTES solution in anhydrous toluene and incubated at room temperature for 2 h. For vinyltrimethoxysilane (VTMS) silanization, substrates were incubated in a 4 % (v/v) VTMS solution in anhydrous toluene for 18 h at room temperature. Silanization with (1-naphthylmethyl)trichlorosilane (NMTS) was conducted according to established protocols described previously[1]. After the respective incubation periods, substrates were thoroughly rinsed sequentially with toluene and ethanol to remove any residues. Finally, the substrates were dried under a continuous nitrogen gas stream and used immediately.

#### Water contact angle measurements

Water contact angles were measured using a Krüss DSA100 drop shape analyzer operating in sessile drop mode. For each measurement, sessile water droplets of 2 µl were deposited on the substrate using a cannula with a diameter of 0.522 mm. Measurements were performed at room temperature (23 °C) and controlled relative humidity (31 %). For each substrate, three to four replicate samples were prepared. On each sample, six measurements of the water contact angle were acquired at different positions, and the average value per sample was calculated. The water contact angle for each substrate is reported as the mean ± standard deviation of the sample average values.

#### Surface-sensitive detection by RIFS-TIRFS

Real-time binding kinetics and surface coverage of functionalized polymers and proteins were characterized using a home-built system for simultaneous reflectance interference spectroscopy (RIFS) and total internal reflection fluorescence spectroscopy (TIRFS), as previously described in detail.[3, 4] Functionalized transducer chips were securely mounted into a flow chamber, equilibrated with buffer, and subsequently exposed to the analyte. Binding events, including polymer adsorption, subsequent protein immobilization, and protein-protein interactions, were monitored under continuous flow conditions. Data acquisition and quantitative extraction of kinetic parameters and binding amplitudes were performed using BIAevaluation 3.1 software (GE Healthcare).[5]

#### Capillary nanostamping

Mesoporous silica stamps were produced according to previously published protocols.[6, 7] Following synthesis, silica monoliths were neutralized and extensively washed with Milli-Q water at 60 °C for 24 h to thoroughly remove residual surfactants and chemical contaminants. Subsequently, the solvent was exchanged to ethanol for another 24 h, during which ethanol was refreshed at least three times to ensure complete solvent replacement. The ethanol-equilibrated stamps were stored for one week before usage. Prior to the stamping procedure, silica stamps were equilibrated in Milli-Q water for 2 hours. Subsequently, stamps were incubated with 20 µL of the respective ink solution for 15 min to allow homogeneous ink absorption into the mesoporous structure. Excess ink was carefully removed using a gentle nitrogen stream. Ink-loaded stamps were mounted onto a stamp holder using double-sided adhesive tape and then gently contacted with silanized substrates for approximately one second to ensure accurate and consistent ink transfer. Following the stamping step, patterned substrates were passivated by incubation with a 1 mg/mL solution of PLL-PEG-OMe for 45 min to minimize nonspecific protein adsorption on non-patterned regions. For cell-based experiments, passivation was performed with a 90:10 (v/v) mixture of PLL-PEG-OMe and PLL-PEG-RGD.[8] Finally, substrates were washed with Milli-Q water and stored refrigerated until experimental applications. For *in vitro* patterning assays, protein concentration and incubation times are given in Table S1.

**Protein expression and purification**

All recombinant fusion proteins utilized in this study were expressed in *Escherichia coli* strain BL21(DE3) Rosetta cells. Gene sequences encoding the respective proteins (Table S1) were cloned into the pET21a vector system, incorporating an N - or C-terminal polyhistidine affinity tag (typically hexahistidine or octahistidine) to facilitate subsequent purification via immobilized metal ion affinity chromatography (IMAC). Bacterial cells were cultured in LB medium supplemented with antibiotics (carbenicillin at 100 µg/mL), and protein expression was induced at an optical density of approximately 0.6 by the addition of 0.5 mM isopropyl-β-D-thiogalactopyranoside (IPTG). Induction proceeded for 4-5 hours at 37 °C. Subsequently, cells were harvested by centrifugation (typically 6000 × g, 15 min, 4 °C). For protein extraction, bacterial pellets were resuspended in ice-cold lysis buffer (20 mM HEPES, pH 7.5, 300 mM NaCl), supplemented with DNase, lysozyme, and EDTA-free protease inhibitor cocktail. Cell disruption was achieved through ultrasonication on ice, followed by clarification of lysates by ultracentrifugation (100000 × g, 45 min, 4 °C). The clarified supernatant was filtered through a 0.45 µm membrane to remove residual particles. Purification was carried out by applying the cleared lysate to a pre-equilibrated HisTrap HP column (GE Healthcare) for IMAC-based affinity chromatography. Protein binding was performed using standard IMAC binding conditions (lysis buffer containing 10 - 20 mM imidazole), followed by extensive washing to remove nonspecifically bound contaminants (lysis buffer supplemented with 20 - 50 mM imidazole). Target proteins were eluted using an increasing gradient of imidazole concentration (typically 250 - 500 mM). Following IMAC purification, protein-containing fractions were pooled and subjected to size exclusion chromatography (SEC) on a HighLoad Superdex 200 column (GE Healthcare), equilibrated with storage buffer (20 mM HEPES, pH 7.5, 150-300 mM NaCl). Fractions corresponding to the monomeric form of the protein (as determined by chromatographic elution profile and SDS-PAGE analysis) were combined, concentrated, if necessary, aliquoted, frozen in liquid nitrogen, and stored at - 80 °C. Purity and integrity of all recombinant proteins were routinely verified by SDS-PAGE, and spectrophotometric quantification prior to application in subsequent experiments.

**Cell culture and transfection**

HeLa cells were cultured in Minimal Essential Medium (MEM) supplemented with Earle's salts and Phenol red, 10 % fetal bovine serum (FBS), 2 mM L-alanyl-L-glutamine, 1 % non-essential amino acids, and 10 mM HEPES buffer. Cells were cultured at 37 °C in a humidified atmosphere containing 5 % $CO_2$. For transient transfection, cells at approximately 70-80 % confluence were detached using a 1× Trypsin-EDTA solution and subsequently seeded into 60-mm tissue culture dishes. After cell attachment, transient transfection was performed using polyethyleneimine (PEI). For each transfection, ~ 5 µg plasmid DNA (detailed plasmid information in Table S2) was diluted in 300 µL of 150 mM NaCl, briefly vortexed, and mixed with 10 µL PEI solution. The DNA-PEI mixture was incubated at room temperature for 15 min before being added dropwise to the culture dishes. Following an incubation period of 5 - 7 hours, cells were washed twice with phosphate-buffered saline (PBS), provided with fresh growth medium, and cultured overnight. After 24 hours, transfected cells were detached by Trypsin-EDTA treatment, reseeded onto pre-functionalized glass coverslips, and incubated for 6 hours before microscopy imaging.

**Binary patterning assay**

Binary patterning was performed using capillary nanostamping with sequential and two-in-one approaches. In sequential binary patterning, substrates underwent two consecutive stamping cycles with two distinct functionalized carrier polymers (PPA combined with PPH or PPS). Slight manual misalignment between stamping cycles resulted in well-defined, spatially offset binary patterns. This facilitated orthogonal protein immobilization, either via distinct transmembrane adaptor proteins or through direct capture of specifically tagged protein fusions. In contrast, the two-in-one binary patterning strategy involved a single stamping step using PPA, subsequently combined with different adaptor proteins (ALFAnb-X). Sequential binary patterning in live-cell experiments employed orthogonal transmembrane adaptor proteins (TA-2m22 and TS-GFPnb) to achieve selective recruitment of cytosolic proteins (tdmCherry and mEGFP) into spatially separated nanodots.

**Total internal reflection fluorescence microscopy (TIRFM)**

TIRF imaging was performed at 25 °C using an inverted microscope (Olympus IX-83) equipped with a modular two-deck configuration. The upper deck was equipped with a 4-line TIRF condenser (cellTIRF (MITICO), Olympus), while the lower deck contained the cellFRAP module (Olympus) for photobleaching experiments. The system was equipped with multiple laser excitation sources, including a 405 nm laser (BCL-100-405, CrystaLaser), a 488 nm diode laser (LuxX 488-200, Omicron), a 561 nm fiber laser (2RU-VFL-P-500-561-B1R, MPB Communications), and a 642 nm fiber laser (2RU-VFL-P-500-642-B1R, MPB Communications). Fluorescence excitation and emission were controlled via a TIRF pentaband polychroic beamsplitter (Semrock zt405/488/561/640/730rpc) and a penta-bandpass emitter filter (BrightLine HC 440/521/607/694/809). Additional single-bandpass filters were used for channel-specific detection: blue (Semrock BrightLine HC 445/45), green (Semrock BrightLine HC 525/35), orange (Chroma 600/50 ET), and red (Chroma 685/50). The system was further equipped with a motorized ultrasonic XY-stage (IX3-SSU, Olympus), and a hardware autofocus system (IX3-ZDC2, 830 nm version, Olympus). Images were acquired with a Hamamatsu ORCA-FusionBT sCMOS camera (2304×2304 pixel resolution) using a 100× oil immersion objective (Olympus UPLAPO 100× HR, NA 1.5). Imaging was controlled via CellSens Dimension software (Version 3.1, Olympus).

For fluorescence recovery after photobleaching (FRAP) experiments on cNDAs, the cellFRAP module (Olympus) was utilized in conjunction with a 405 nm bleaching laser (LuxX+ 405-60, Omicron). Image acquisition was conducted using a three-line polychroic beamsplitter (Chroma, zt488/561/633/405tpc) located in the upper deck, while the lower deck incorporated a FRAP cube consisting of a dichroic beamsplitter at 405 nm (Chroma, H 405 LPXR superflat) and a QuadLine emitter filter (Chroma, zet405/488/561/640m).

**Fluorescence recovery after photobleaching (FRAP)**

FRAP experiments were conducted by initially acquiring three pre-bleach fluorescence images of a defined circular region of interest (ROI) with a diameter of approximately 15 µm. Following photobleaching, fluorescence recovery within this region was monitored continuously at frame rates of either 0.1 frame per second or 1 frame per second over a total duration of 3 min. To quantify recovery kinetics, individual nanodots (typically 20 - 40 per region) were selected within the bleached ROI, and fluorescence intensities were measured and analyzed using a custom-written MATLAB script.

FRAP curves were calculated according to the following equation[9]:

$$F(t)=\frac{\left(F_{ROI_{inside}}\text{-}F_{offset}\right)\text{-}\left(F_{ROI_{outside}}\text{-}F_{offset}\right)}{\left(\frac{F_{ref}\text{-}F_{offset}}{F_{\mathrm{re}f_0}\text{-}F_{offset}}\right)}$$

Here, $F_{ROI_{inside}}$and $F_{ROI_{outside}}$denote the measured fluorescence intensities inside and outside the patterned areas, respectively, within the bleached region. $F_{ref}$ is the fluorescence intensity of an unbleached reference region within the pattern, while $F_{\mathrm{re}f_0}$ represents the fluorescence intensity of the same reference region prior to bleaching. The offset intensity $F_{offset}$ was determined from a region outside of the cells and subsequently subtracted from all intensity measurements. Fluorescence recovery curves were fitted using a monoexponential decay function to characterize recovery kinetics. All data processing, normalization, and visualization were performed using MATLAB and OriginPro 9.

**DNA-PAINT imaging and analysis**

DNA-PAINT experiments were performed on samples prepared *in vitro* as well as on fixed cellular samples. For *in vitro* experiments, patterned substrates were stained with 20 nM anti-tag nanobody (MASSIVE-TAG-X2-FAST anti-GFP or MASSIVE-TAG-Q-FAST anti-ALFA) coupled to the DNA docking strand 3 or F3 for 20 min at room temperature. Excess was removed by extensive washing steps with washing buffer. Cellular samples were fixed using 4 % paraformaldehyde (PFA) for 15 min at room temperature, washed five times with PBS, permeabilized with 0.1 % (v/v) Triton-X100 in PBS for 10 min, and blocked with 3 % (w/v) BSA in PBS for 30 min. After removing the blocking solution, samples were incubated overnight at 4 °C with 50 nM anti-GFP nanobody conjugated to docking strand F3. Subsequently, the samples were washed thoroughly and incubated with 90 nm gold nanoparticles as fiducial markers (Cytodiagnostics, G-90-20, 1:1000 dilution) for 15 min, followed by additional washing steps. DNA-PAINT imaging was performed using the previously described TIRF microscope setup. Samples were incubated in imaging buffer containing 250-500 pM Cy3B-conjugated imager strand (500 pM for *in vitro* experiments). Imaging was conducted at 25 °C with a 561 nm excitation laser adjusted to a power density of approx. 100 W/cm$^2$ in the focal plane. To avoid spectral crosstalk between tdmCherry and the Cy3B imager strands, residual tdmCherry fluorescence was photobleached in the 561 nm channel for 60 s at imaging conditions immediately before starting DNA-PAINT acquisition. Typically, datasets consisted of 40000 frames recorded at 50 ms or 100 ms exposure time with a 2×2 pixel binning (effective pixel size: 130 nm). A hardware autofocus system ensured stable focus throughout the acquisition. Raw datasets were analyzed using the Picasso software suite.[10] Single-molecule localizations were identified using Picasso Localize with a box size of 7 pixels and a minimum net gradient threshold set between 5000-20000 to exclude low-quality localizations. Photon conversion parameters were set as follows: EM gain: 1; baseline: 400; sensitivity: 0.26; quantum efficiency: 0.92; pixel size: 130 nm. Localizations were fitted with an integrated Gaussian function[11] and subsequently filtered in Picasso Filter for localization precision ≤ 6.5 nm (0.05 pixels). Drift correction was initially performed by cross-correlation followed by refinement using fiducial markers. Representative averaged images from *in vitro* experiments were generated using Picasso Average, aligning and averaging multiple nanodots (>200 per dataset). Parameters were set as follows: oversampling: 40, iterations: 10. Quantitative parameters for *in vitro* experiments - including nanodot diameter, localization density, peak intensity, and contrast - were determined from DNA-PAINT datasets. Diameters were measured manually based on localization boundaries. Localization density was quantified per defined nanodot area (calculated from measured diameters), peak intensities were measured as the maximum localization count per unit area (130 nm pixel size), and contrast was calculated as the ratio of peak intensity to local background intensity. DBSCAN cluster analysis was employed to identify protein clusters by grouping localizations based on spatial density without assuming any predefined cluster shape. The analysis was performed using a custom-written MATLAB script with a minimum localization threshold of 22 and a radius of 7.5 nm. Extracted parameters included the percentage of clustered localizations per nanodot, total localizations per cluster, and cluster diameter.

**RESI imaging and analysis**

RESI imaging was performed on fixed cell samples prepared analogously to DNA-PAINT protocols.[12] Cells expressing GFP-tagged proteins (IRAK4-mEGFP) were fixed with 4 % PFA for 15 min at room temperature, washed thoroughly with PBS, permeabilized with 0.1 % Triton-X100 in PBS for 10 min, and blocked with 3 % BSA in PBS for 30 min. After blocking, samples were incubated overnight at 4 °C with anti-GFP single-domain antibodies (sdAB) conjugated to four distinct RESI docking strands, each at an equal concentration of 25 nM (total concentration: 100 nM). After washing, samples were incubated with fiducial markers for 15 min, followed by further washing. Sequential RESI imaging rounds were conducted using Cy3B-conjugated imager strands ($I_1$-$I_4$), each complementary to one of the four docking strands. Each imaging round consisted of incubating samples with 1 nM of a single imager strand in imaging buffer, typically acquiring 20000 frames at 50 ms exposure time with 2×2 pixel binning (effective pixel size: 130 nm). Samples were excited with a 561 nm laser adjusted to a power density of approx. 100 W/cm$^2$ in the focal plane, and stable imaging conditions were maintained using a hardware autofocus. To avoid spectral crosstalk between tdmCherry and the Cy3B imager strands, residual tdmCherry fluorescence was photobleached in the 561 nm channel for 60 s at imaging conditions immediately before starting RESI imaging. Raw RESI datasets were processed individually using the Picasso software suite.[10] Single-molecule localizations were identified using Picasso Localize with a box size of 7 pixels and a minimum net gradient threshold set between 5000 and 10000. Photon conversion parameters were identical to those described for DNA-PAINT. Localizations were fitted with an integrated Gaussian function and filtered for localization precision ≤ 5.2 nm. Drift correction between sequential imaging rounds was initially performed via cross-correlation and further refined using fiducial markers to ensure precise channel alignment. RESI post-processing was performed in Picasso Render applying a cluster radius of 10 nm and a minimum localization threshold of 10 localizations. Further quantitative analysis was performed using a custom-written MATLAB script. RESI localizations were imported, and an interaction radius (9.5 nm) was chosen based on structural considerations from protein dimensions. Molecular clusters were automatically identified by grouping localizations whose pairwise distances were within twice this interaction radius. These clusters were classified as monomers, dimers, trimers, tetramers, or higher-order oligomers according to the number of RESI localizations per group. Spatial metrics, including pairwise intermolecular distances, were determined for each cluster. Additionally, quantitative parameters were directly obtained from Picasso Render: Number of molecules per nanodot, RESI localization density per nanodot, and localization precision per individual molecule.

**Image evaluation**

Fluorescence intensity line profiles were generated from single-, dual-, or multi-color TIRF microscopy images using Fiji (ImageJ, Version 1.54f).[13] Line profiles were manually positioned across representative nanodots to qualitatively evaluate protein co-localization and spatial overlap. Quantitative fluorescence intensity analyses of individual nanodots were performed using a custom-written MATLAB script. Nanodots were manually selected from multiple images, and peak fluorescence intensities were quantified as the maximum values derived from intensity line profiles. Contrast values were calculated as the ratio between peak fluorescence intensity ($I_{max}$) and the adjacent local minimum background intensity ($I_{min}$).

$$Contrast = \frac{I_{max} - I_{min}}{I_{min}}$$

Quantitative co-localization analysis in dual- or multi-color TIRF microscopy images was determined using Pearson's correlation coefficients (PCC). PCC values were calculated from multiple defined regions of interest (ROIs) per image and are presented as mean ± standard deviation (SD). Single-cell analyses of contrast and peak intensity were conducted by quantifying multiple nanodots within single cells. The correlation between immobilized proteins (bait) and co-recruited proteins (prey) was assessed through change-point threshold analysis and linear regression using MATLAB and OriginPro 9. Relative contrast and relative peak intensity values were calculated by dividing fluorescence intensities of prey proteins by the corresponding immobilized bait proteins, enabling direct comparisons across experimental conditions. For measurements involving PCC, peak intensities, contrast, and relative values, 5 - 10 cells per condition were analyzed, with a total of > 1000 nanodots evaluated. Statistical significance between experimental groups was assessed using Kolmogorov-Smirnov and Tukey tests. Data are represented as mean ± SD, unless otherwise stated, with statistical significance indicated as follows: *p ≤ 0.05, **p ≤ 0.01, ***p ≤ 0.001, ****p ≤ 0.0001. Image processing, quantification, and statistical analyses were performed using Fiji, MATLAB (R2023a), and OriginPro 9 (OriginLab Corporation).

**Structural Prediction using AlphaFold3**

Structural predictions of the myddosome complex were generated using the AlphaFold3 online server. Protein sequences for human MyD88 (hMyD88; residues M54-I109), mouse IRAK4 (mIRAK4; residues M1-A460), and mouse IRAK1 (mIRAK1; residues M1-F688) were retrieved from the UniProt database. To model the hierarchical assembly of the myddosome, multimeric complexes comprising six copies of hMyD88, four copies of mIRAK4, and four copies of mIRAK1 were predicted. AlphaFold3 generated five structural models per multimeric complex, applying 10 recycling iterations to ensure structural refinement.[14] For detailed investigation of the core oligomerization interface, additional predictions were performed using only the death domain (DD) regions of hMyD88 (M54-I109), mIRAK4 (R20-A104), and mIRAK1 (M27-A106). Structural visualization and analysis were conducted using ChimeraX 1.8.

### Supplementary References

(1) Philippi, M.; Richter, C. P.; Kappen, M.; Watrinet, I.; Miao, Y.; Runge, M.; Jorde, L.; Korneev, S.; Holtmannspotter, M.; Kurre, R.; et al. Biofunctional Nanodot Arrays in Living Cells Uncover Synergistic Co-Condensation of Wnt Signalodroplets. *Small* **2022**, *18* (50), e2203723. DOI: 10.1002/smll.202203723.
(2) Wedeking, T.; Lochte, S.; Birkholz, O.; Wallenstein, A.; Trahe, J.; Klingauf, J.; Piehler, J.; You, C. Spatiotemporally Controlled Reorganization of Signaling Complexes in the Plasma Membrane of Living Cells. *Small* **2015**, *11* (44), 5912-5918. DOI: 10.1002/smll.201502132.
(3) Gavutis, M.; Lata, S.; Piehler, J. Probing 2-dimensional protein-protein interactions on model membranes. *Nat Protoc* **2006**, *1* (4), 2091-2103. DOI: 10.1038/nprot.2006.270.
(4) Gavutis, M.; Lata, S.; Lamken, P.; Muller, P.; Piehler, J. Lateral ligand-receptor interactions on membranes probed by simultaneous fluorescence-interference detection. *Biophys J* **2005**, *88* (6), 4289-4302. DOI: 10.1529/biophysj.104.055855.
(5) Bhagawati, M.; You, C.; Piehler, J. Quantitative real-time imaging of protein-protein interactions by LSPR detection with micropatterned gold nanoparticles. *Anal Chem* **2013**, *85* (20), 9564-9571. DOI: 10.1021/ac401673e.
(6) Schmidt, M.; Philippi, M.; Münzner, M.; Stangl, J. M.; Wieczorek, R.; Harneit, W.; Müller - Buschbaum, K.; Enke, D.; Steinhart, M. Capillary Nanostamping with Spongy Mesoporous Silica Stamps. *Advanced Functional Materials* **2018**, *28* (23). DOI: 10.1002/adfm.201800700.
(7) Philippi, M.; You, C.; Richter, C. P.; Schmidt, M.; Thien, J.; Lisse, D.; Wollschlager, J.; Piehler, J.; Steinhart, M. Close-packed silane nanodot arrays by capillary nanostamping coupled with heterocyclic silane ring opening. *RSC Adv* **2019**, *9* (43), 24742-24750. DOI: 10.1039/c9ra03440d From NLM PubMed-not-MEDLINE.
(8) VandeVondele, S.; Voros, J.; Hubbell, J. A. RGD-grafted poly-L-lysine-graft-(polyethylene glycol) copolymers block non-specific protein adsorption while promoting cell adhesion. *Biotechnol Bioeng* **2003**, *82* (7), 784-790. DOI: 10.1002/bit.10625.
(9) Lochte, S.; Waichman, S.; Beutel, O.; You, C.; Piehler, J. Live cell micropatterning reveals the dynamics of signaling complexes at the plasma membrane. *The Journal of cell biology* **2014**, *207* (3), 407-418. DOI: 10.1083/jcb.201406032.
(10) Schnitzbauer, J.; Strauss, M. T.; Schlichthaerle, T.; Schueder, F.; Jungmann, R. Super-resolution microscopy with DNA-PAINT. *Nature Protocols* **2017**, *12* (6), 1198-1228. DOI: 10.1038/nprot.2017.024.
(11) Mortensen, K. I.; Churchman, L. S.; Spudich, J. A.; Flyvbjerg, H. Optimized localization analysis for single-molecule tracking and super-resolution microscopy. *Nature Methods* **2010**, *7* (5), 377-381. DOI: 10.1038/nmeth.1447.
(12) Reinhardt, S. C. M.; Masullo, L. A.; Baudrexel, I.; Steen, P. R.; Kowalewski, R.; Eklund, A. S.; Strauss, S.; Unterauer, E. M.; Schlichthaerle, T.; Strauss, M. T.; et al. Ångström-resolution fluorescence microscopy. *Nature* **2023**, *617* (7962), 711-716. DOI: 10.1038/s41586-023-05925-9.
(13) Schindelin, J.; Arganda-Carreras, I.; Frise, E.; Kaynig, V.; Longair, M.; Pietzsch, T.; Preibisch, S.; Rueden, C.; Saalfeld, S.; Schmid, B.; et al. Fiji: an open-source platform for biological-image analysis. *Nature Methods* **2012**, *9* (7), 676-682. DOI: 10.1038/nmeth.2019.
(14) Abramson, J.; Adler, J.; Dunger, J.; Evans, R.; Green, T.; Pritzel, A.; Ronneberger, O.; Willmore, L.; Ballard, A. J.; Bambrick, J.; et al. Accurate structure prediction of biomolecular interactions with AlphaFold 3. *Nature* **2024**, *630* (8016), 493-500. DOI: 10.1038/s41586-024-07487-w.

### Supplementary Figures

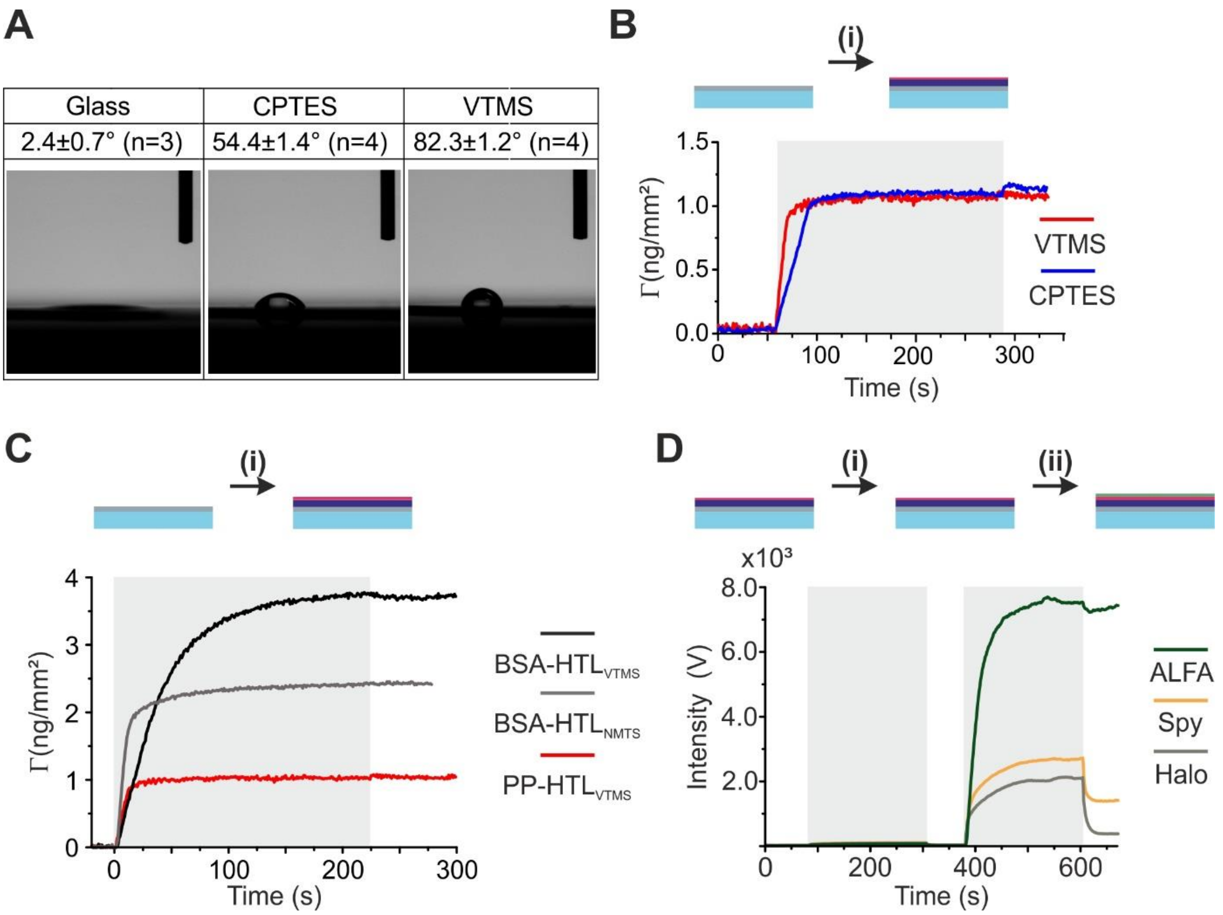


**Figure S1. Characterization of surface silanization and carrier biopolymer interactions.** (A) Representative water contact angle measurements of glass surfaces silanized with VTMS or CPTES, in comparison with plasma-cleaned glass. Results for each substrate are based on measurements from 3 to 4 independent samples (n = 3 or 4). Values are reported as mean ± s.d. of the sample averages. (B) Reflectance interference spectroscopy (RIFS) detection of PPH binding on VTMS- and CPTES-silanized surfaces. (C) Comparison of surface coating kinetics for BSA-HTL on VTMS and NMTS as well as PPH on VTMS. (D) Total internal reflection fluorescence spectroscopy (TIRFS) detection of differently functionalized PP derivatives. mEGFP was injected as a negative control (i), followed by the respective binder fused to mEGFP (ii).

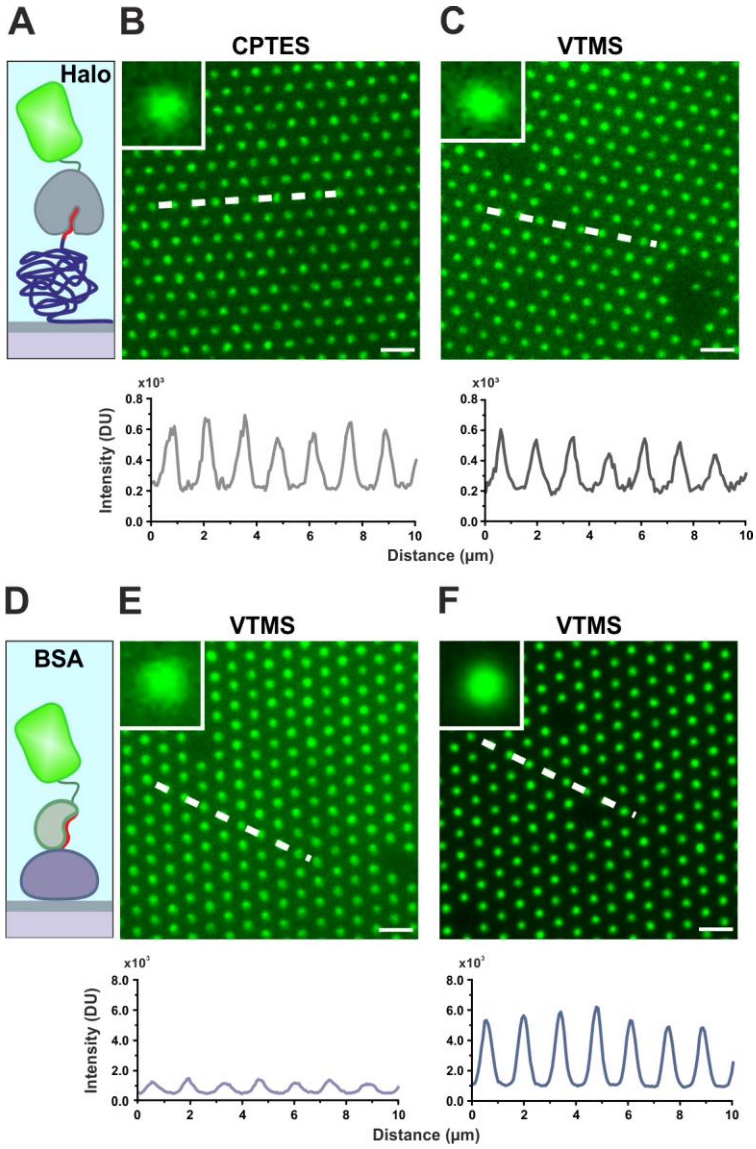


**Figure S2. bNDAs utilizing PPH and BSA.** (A-C) Schematic illustration of PPH/Halo-mEGFP (A), alongside TIRF microscopy images on CPTES (B) and VTMS (C) surfaces. White dashed lines denote regions selected for intensity profiles. Scale bars: 2 µm. Insets provide magnified views of individual nanodots. (D-F) Schematic depiction of BSA-based nanodot arrays (D), with corresponding TIRF microscopy images of bNDAs using BSA-HTL/Halo-mEGFP (E) and BSA-ALFAtag/ALFAnb-mEGFP (F). White dashed lines indicate regions selected for intensity profiles. Scale bars: 2 µm. Insets provide magnified views of individual nanodots.

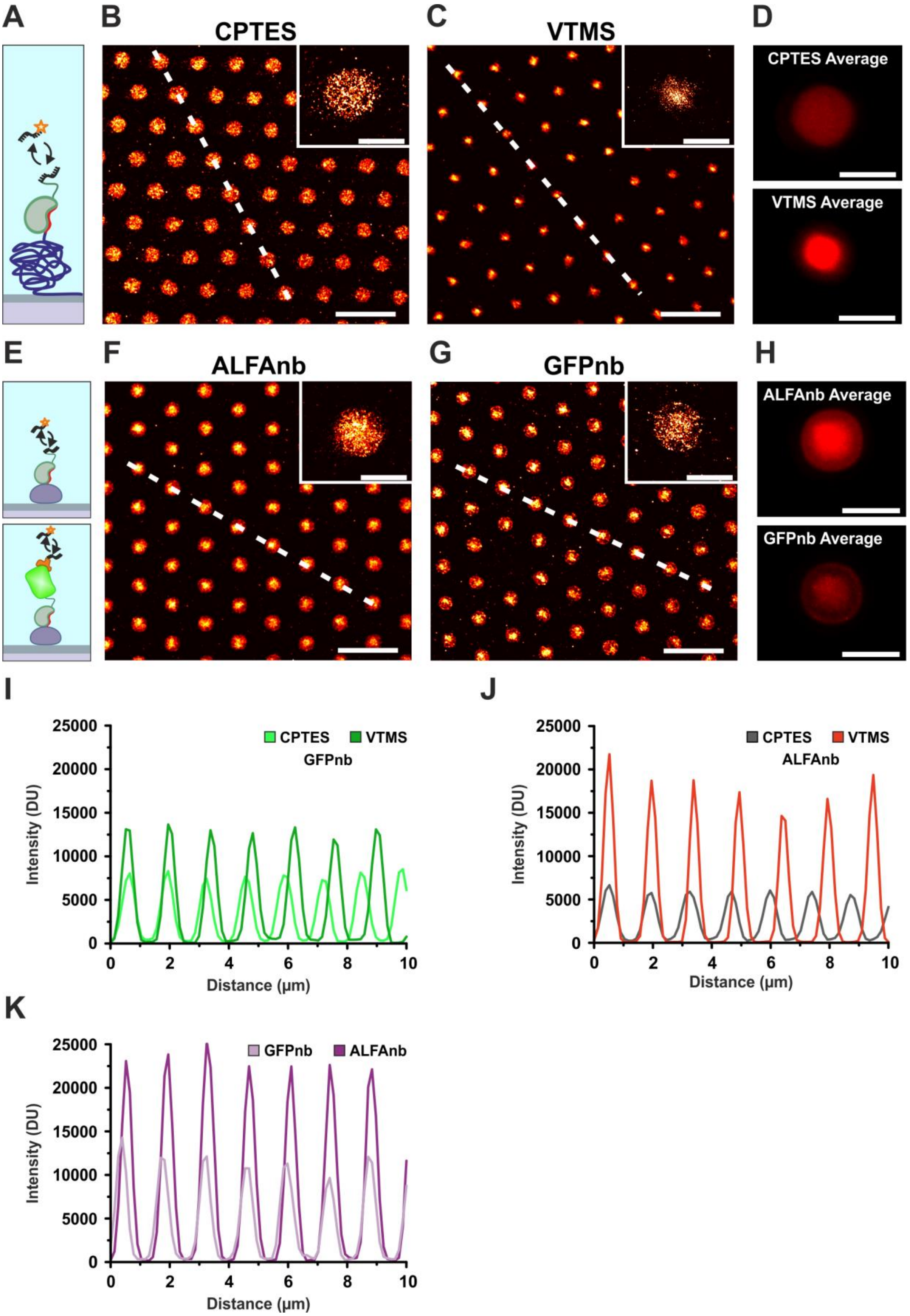


**Figure S3. Nanoscale imaging of PPA and BSA-ALFAtag bNDAs by DNA-PAINT.** (A) Schematic overview of the DNA-PAINT labeling strategy applied to PPA NDAs, utilizing an anti-ALFA nanobody for selective imaging. (B, C) Representative DNA-PAINT super-resolution images obtained on CPTES (B) and VTMS (C) surfaces. White dashed lines indicate areas used for intensity profiles. Scale bars: 2 µm. Insets provide enlarged views of individual nanodots. Scale bars: 500 nm. (D) Averaged DNA-PAINT images generated by aligning and averaging 200 individual nanodots. Scale bars: 500 nm. (E) Schematic depiction of the labeling strategy for BSA-ALFAtag nanodots, employing either anti-ALFA nanobody (top) or anti-GFP nanobody (bottom). (F, G) Representative DNA-PAINT images of nanodots labeled with anti-ALFA (F) or anti-GFP (G) nanobodies. White dashed lines denote regions analyzed for intensity profiles. Scale bars: 2 µm. Insets provide magnified views of individual nanodots. Scale bars: 500 nm. (H) Averaged DNA-PAINT images generated by aligning and averaging 200 individual nanodots. Scale bars: 500 nm. (I-K) Intensity line profiles corresponding to PPA/ALFAnb-mEGFP/GFPnb (I), PPA/ALFAnb (J) and BSA-ALFAtag (K) bNDAs.

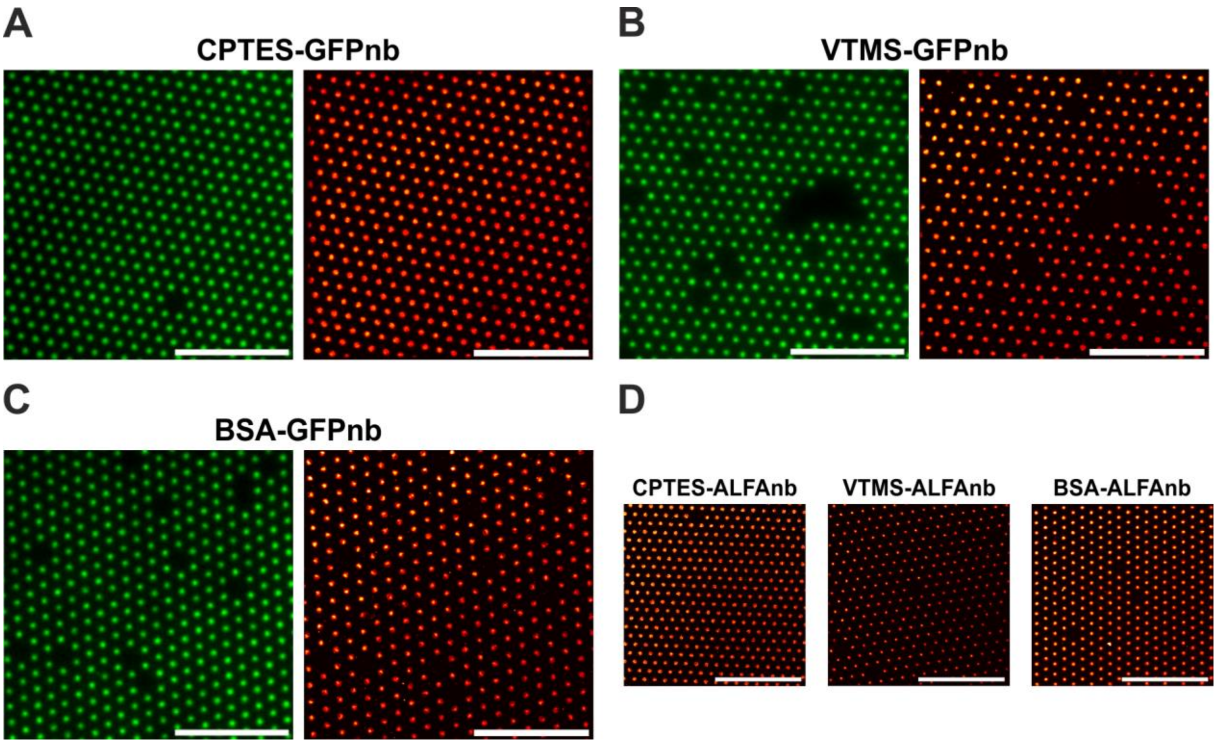


**Figure S4. Full field-of-view TIRF and DNA-PAINT images.** (A-C) TIRF (left) and DNA-PAINT (right) images for PPA bNDAs stained with anti-GFP nanobody on CPTES (A) and VTMS (B) surfaces, as well as for BSA-ALFAtag bNDAs labeled with anti-GFP nanobody (C). (D) DNA-PAINT images of PPA bNDAs labeled with anti-ALFA nanobody on different surfaces (left, middle), alongside corresponding BSA-ALFAtag bNDAs (right). Scale bars: 10 µm.

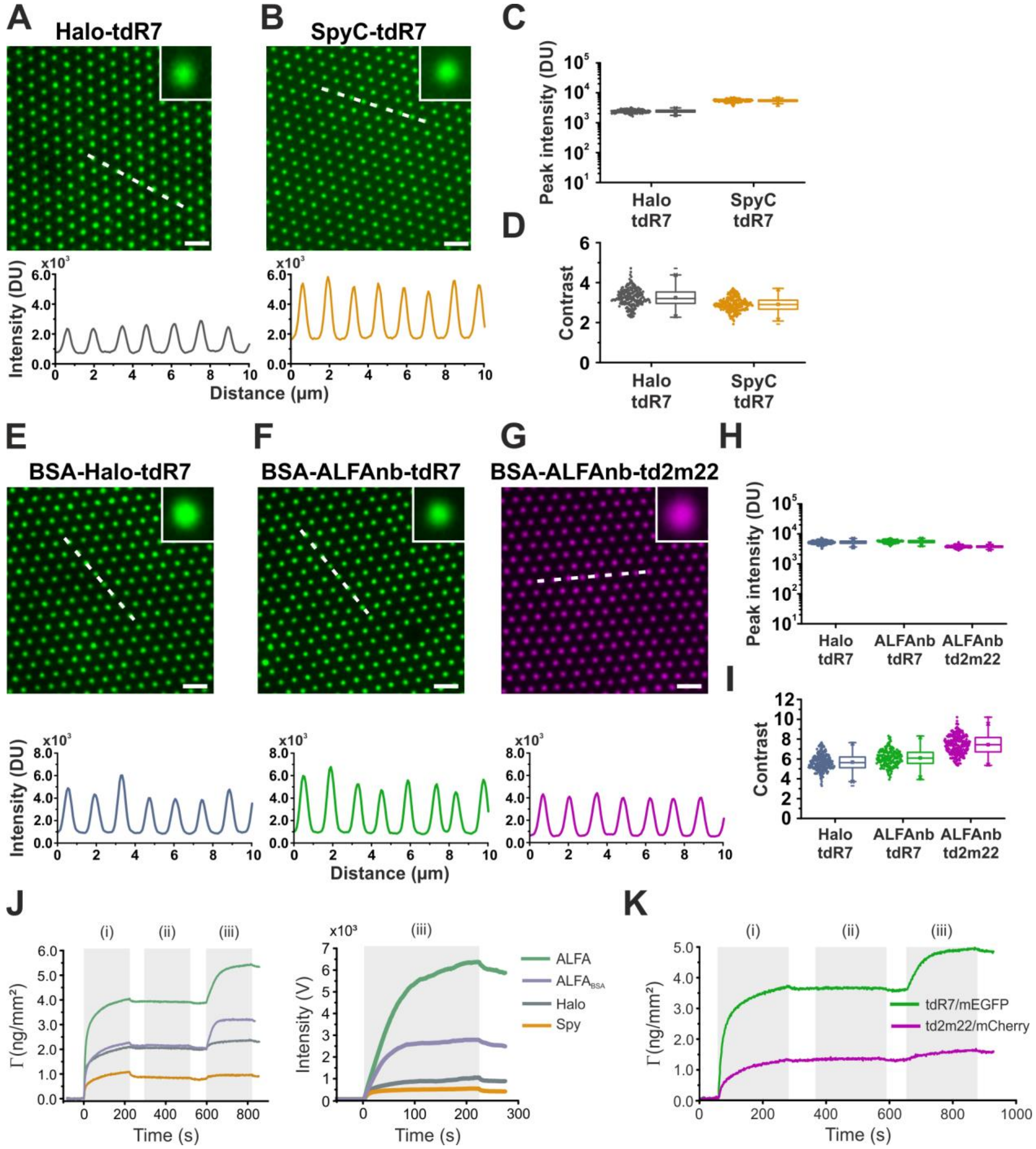


**Figure S5. Characterization of orthogonal tandem adaptor proteins.** (A, B) Representative TIRF microscopy images of Halo-tdR7/mEGFP (A) and SpyCatcher-tdR7/mEGFP (B) bNDAs. Insets provide magnified views of individual nanodots, with white dashed lines indicating regions analyzed for intensity profiles. Scale bars: 2 µm. (C, D) Quantitative comparison of peak fluorescence intensity (C) and contrast (D) for Halo-tdR7/mEGFP and SpyCatcher-tdR7/mEGFP. (E-G) Representative TIRF microscopy images of BSA conjugates (HTL and ALFAtag) using Halo-tdR7/mEGFP (E), ALFAnb-tdR7/mEGFP (F) and ALFAnb-td2m22/mCherry (G) with corresponding intensity profiles. Insets provide magnified views of individual nanodots, with white dashed lines indicating regions analyzed for intensity profiles. Scale bars: 2 µm. (H, I) Quantitative analysis of peak fluorescence intensity (H) and contrast (I) for each respective binding pair. (J) RIFS/TIRFS-based detection of tandem adaptor proteins on PP- and BSA-functionalized surfaces (i). MBP was used as negative control (ii), followed by mEGFP (iii). (K) Orthogonal tandem adaptor binding analysis via RIFS, showing stepwise injection of ALFAnb-tdR7 (green) or ALFAnb-td2m22 (magenta) (i), followed by (ii, iii) sequential binding of mEGFP and mCherry.

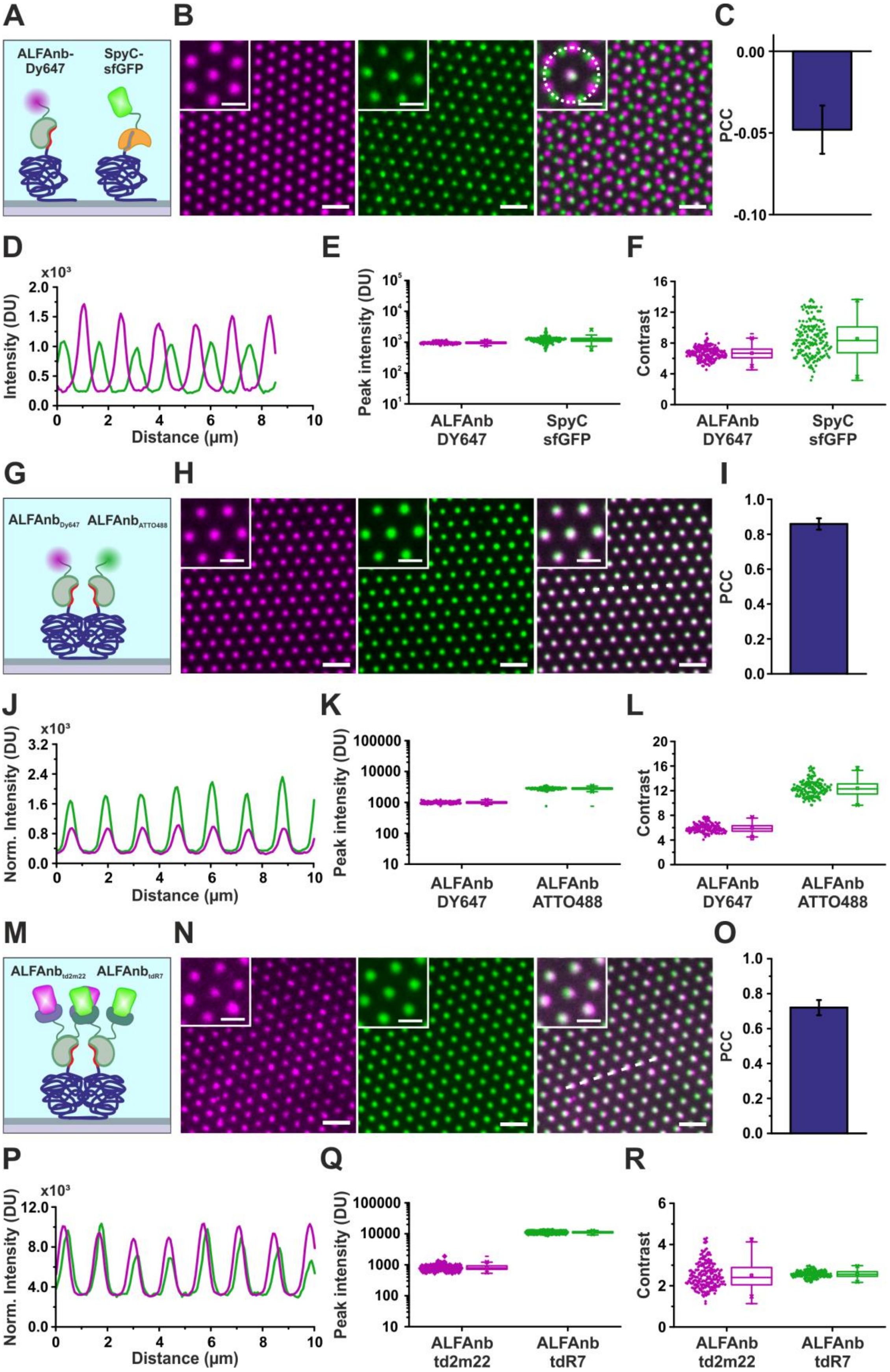


**Figure S6. Binary patterning strategies to control protein co-organization at different scales.** (A) Schematic illustration of the multiplexed biofunctionalization approach combining PPA/ALFAnb-Dy647 and PPS/SpyCatcher-sfGFP. (B) Representative dual-color TIRF microscopy images demonstrating precisely defined binary patterning (right) and distinct immobilization of individual binders (left, middle). Insets display detailed views of individual channels and the composite merged image. Scale bars: 2 µm; insets: 1 µm. (C) PCC analysis quantifying co-localization within the binary patterning framework. (D) Fluorescence intensity line profiles corresponding to (B), highlighting periodic signal alternation. (E, F) Quantitative analysis of fluorescence peak intensities (E) and contrast (F) for the respective channels. (G) Schematic illustration of the multiplexed two-in-one biofunctionalization strategy employing PPA with ALFAnb-Dy647 and ALFAnb-ATTO488. (H) Representative dual-color TIRF microscopy images illustrating two-in-one binary patterning (right) alongside individual fluorescence channels (left, middle). Insets provide detailed views of individual channels and the composite merged image. Scale bars: 2 µm; insets: 1 µm. (I) PCC analysis assessing co-localization within the two-in-one binary patterning system. (J) Fluorescence intensity line profiles corresponding to (H), showing co-localized peaks. (K, L) Quantitative analysis of fluorescence peak intensities (K) and contrast (L) for respective binders. (M) Schematic illustration of the multiplexed two-in-one biofunctionalization strategy incorporating PPA/ALFAnb-td2m22/mCherry and PPA/ALFAnb-tdR7/mEGFP. (N) Representative dual-color TIRF microscopy images depicting two-in-one binary patterning (right) along with single-channel visualizations (left, middle). Insets provide magnified views of individual channels and the composite merged image. Scale bars: 2 µm; insets: 1 µm. (O) PCC analysis quantifying co-localization within the two-in-one binary patterning system. (P) Fluorescence intensity line profiles corresponding to (N), showing co-localized peaks. (Q, R) Quantitative analysis of fluorescence peak intensities (Q) and contrast (R) for the respective binder pairs.

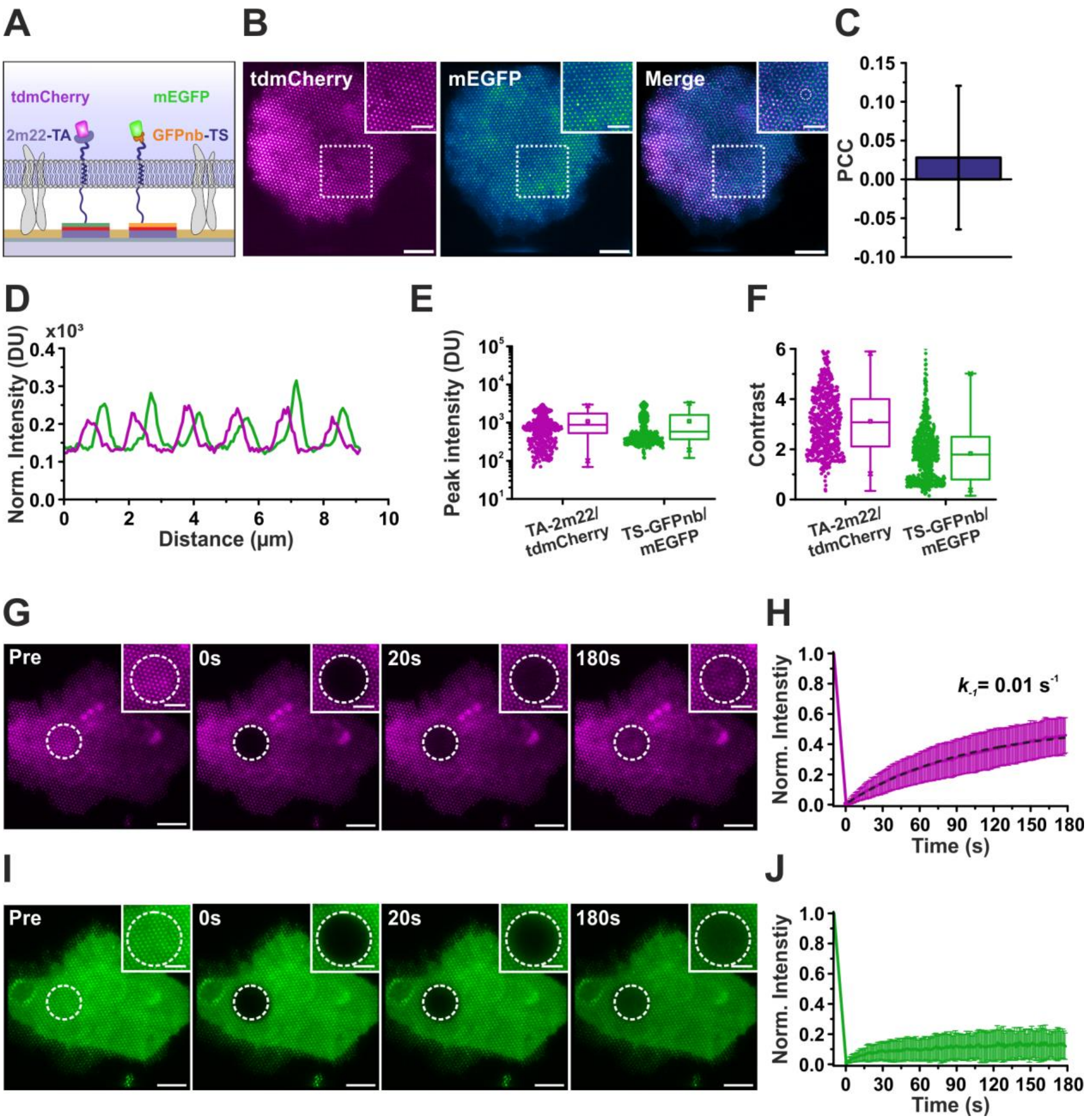


**Figure S7. Binary patterning of transmembrane adaptor proteins in live cells.** (A) Schematic illustration of the multiplexed biofunctionalization strategy integrating PPA/TA-2m22 and PPS/TS-GFPnb for the targeted recruitment of cytosolic tdmCherry and mEGFP, respectively. (B) Representative dual-color TIRF microscopy images displaying the discrete immobilization of tdmCherry (left) and mEGFP (middle) within their respective cNDAs. The merged overlay (right) reveals the precise binary spatial arrangement. Insets provide magnified views of individual nanodots. Scale bars: 10 μm; insets: 5 μm. (C) PCC analysis quantifying co-localization within the binary patterning framework, validating the defined spatial segregation of the immobilized proteins. (D) Fluorescence intensity line profiles corresponding to (B), highlighting alternating intensity peaks. (E, F) Quantitative analysis of peak fluorescence intensity (E) and contrast (F) for TA-2m22/tdmCherry and TS-GFPnb/mEGFP. (G-J) Time-lapse TIRF microscopy capturing FRAP of tdmCherry (G) and mEGFP (I). Dashed circles indicate bleached regions. Scale bars: 10 μm; insets: 5 μm. The corresponding normalized fluorescence intensity recovery curves in (H) and (J) highlight dynamic turnover for tdmCherry and stable retention for mEGFP. The recovery curve for 2m22/tdmCherry was fitted with a monoexponential function (black dashed line) to quantify the dissociation rate constant ($k_{-1}$).

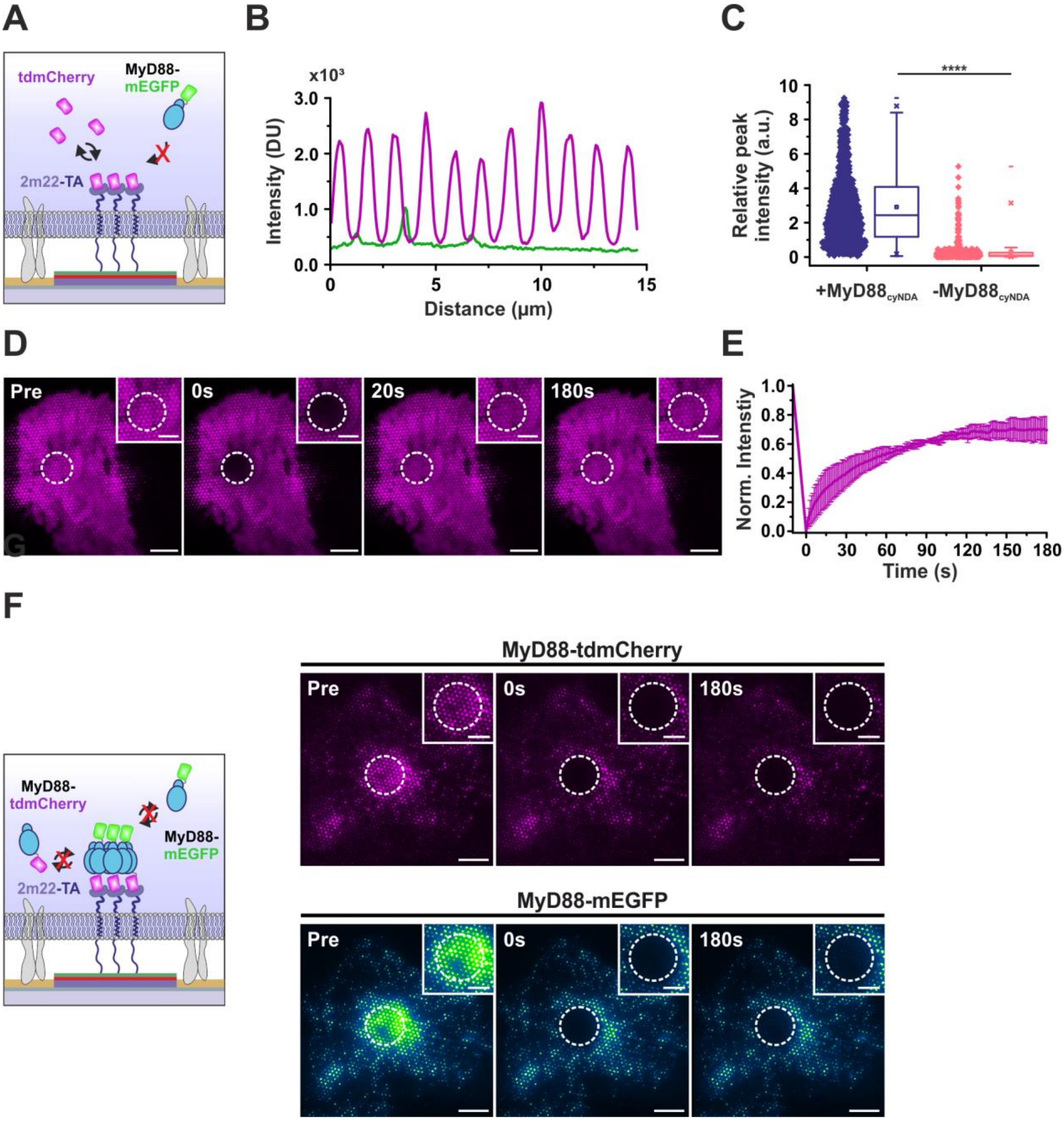


**Figure S8.** **FRAP negative control for tdmCherry in the presence of MyD88-mEGFP.** (A) Schematic illustration depicting the immobilization of tdmCherry via TA-2m22 adaptor transmembrane protein in the presence of cytosolic MyD88-mEGFP. (B) Fluorescence intensity line profiles for tdmCherry (magenta) and MyD88-mEGFP (green). (C) Quantitative comparison of relative peak intensities in the presence (+) and absence (-) of immobilized MyD88-tdmCherry within cNDAs (****$p < 0.0001$). (D) Time-lapse TIRF microscopy capturing FRAP of tdmCherry (D) in the presence of MyD88-mEGFP. Dashed circles indicate bleached regions. Insets highlight individual nanodots Scale bars: 10 µm; insets: 5 µm. (E) Normalized fluorescence recovery curve for tdmCherry showing the dynamic turnover in the presence of MyD88-mEGFP. (F) Schematic illustration and representative dual-color FRAP images of MyD88-tdmCherry (magenta, top) and MyD88-mEGFP (green, bottom), showing the absence of fluorescence recovery, indicative of stable protein complexes formed by MyD88 oligomerization. Dashed circles indicate bleached regions. Scale bars: 10 µm; insets: 5 µm.

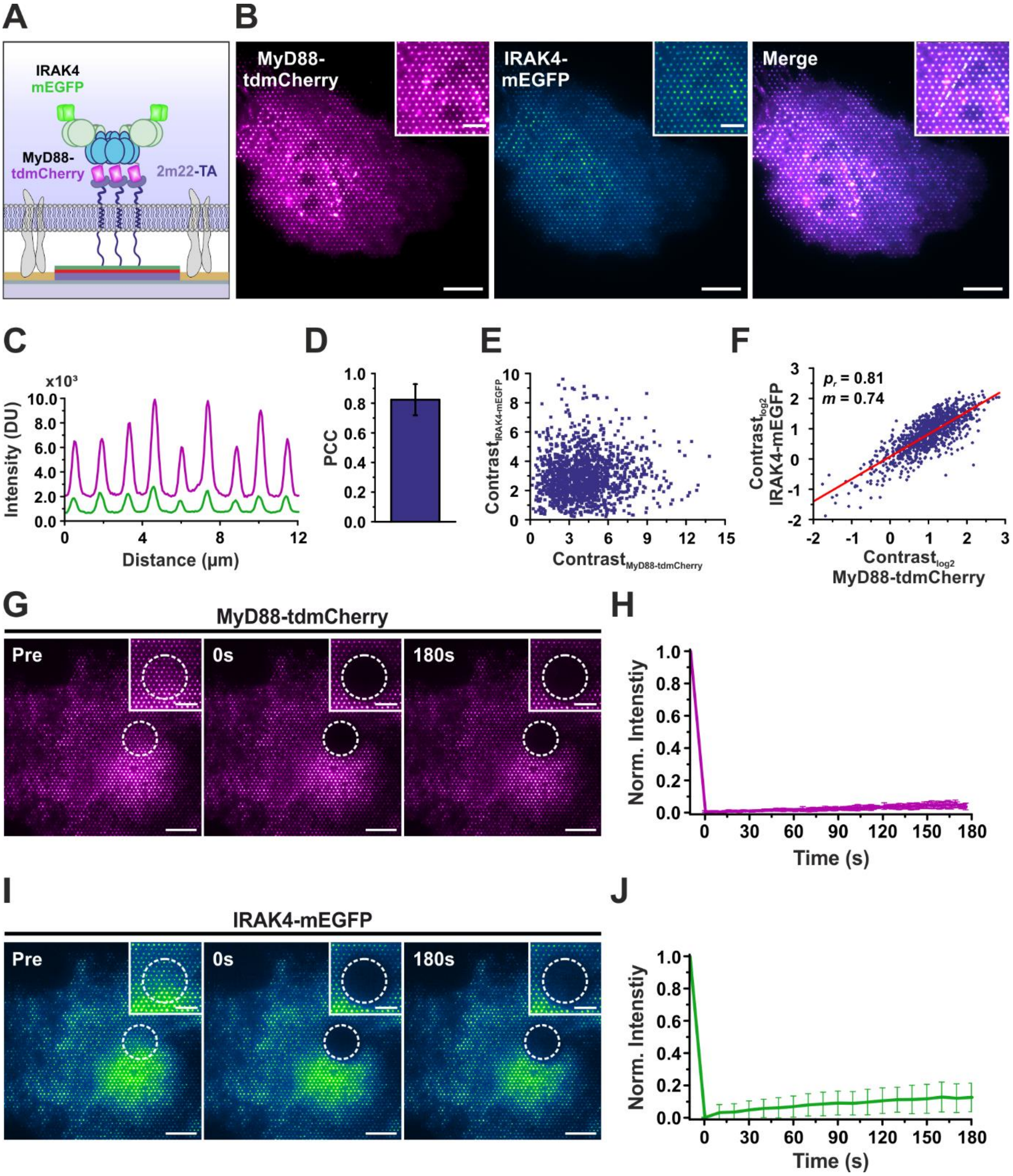


**Figure S9. Co-recruitment of IRAK4 to immobilized MyD88 complexes and complex stability analysis by FRAP.** (A) Schematic representation depicting the immobilization of MyD88-tdmCherry within cNDAs mediated by the TA-2m22 adaptor transmembrane protein, facilitating the targeted co-recruitment of cytosolic IRAK4-mEGFP. (B) Representative dual-color TIRF microscopy images showing immobilized MyD88-tdmCherry (magenta, left), co-recruited IRAK4-mEGFP (green, middle), and merged overlay (right). Insets provide magnified views of individual nanodots. Scale bars: 10 µm; insets: 5 µm. (C) Fluorescence intensity line profiles derived from (B) demonstrate overlapping peaks of MyD88-tdmCherry and IRAK4-mEGFP, confirming co-recruitment within cNDAs. (D) PCC quantifying the degree of co-localization between MyD88-tdmCherry and IRAK4-mEGFP. (E) Scatter plot depicting contrast values for individual nanodots, providing insight into co-recruitment efficiency. (F) Single-cell correlation analysis between contrast values of MyD88-tdmCherry and IRAK4-mEGFP for approximately 1000 individual nanodots, highlighting the linear co-recruitment behavior

(Pearson's r-value and slope indicated). (G-J) Time-lapse TIRF microscopy images and corresponding FRAP recovery curves for MyD88-tdmCherry (G, H) and IRAK4-mEGFP (I, J). Dashed circles indicate bleached regions. Scale bars: 10 µm; insets: 5 µm.

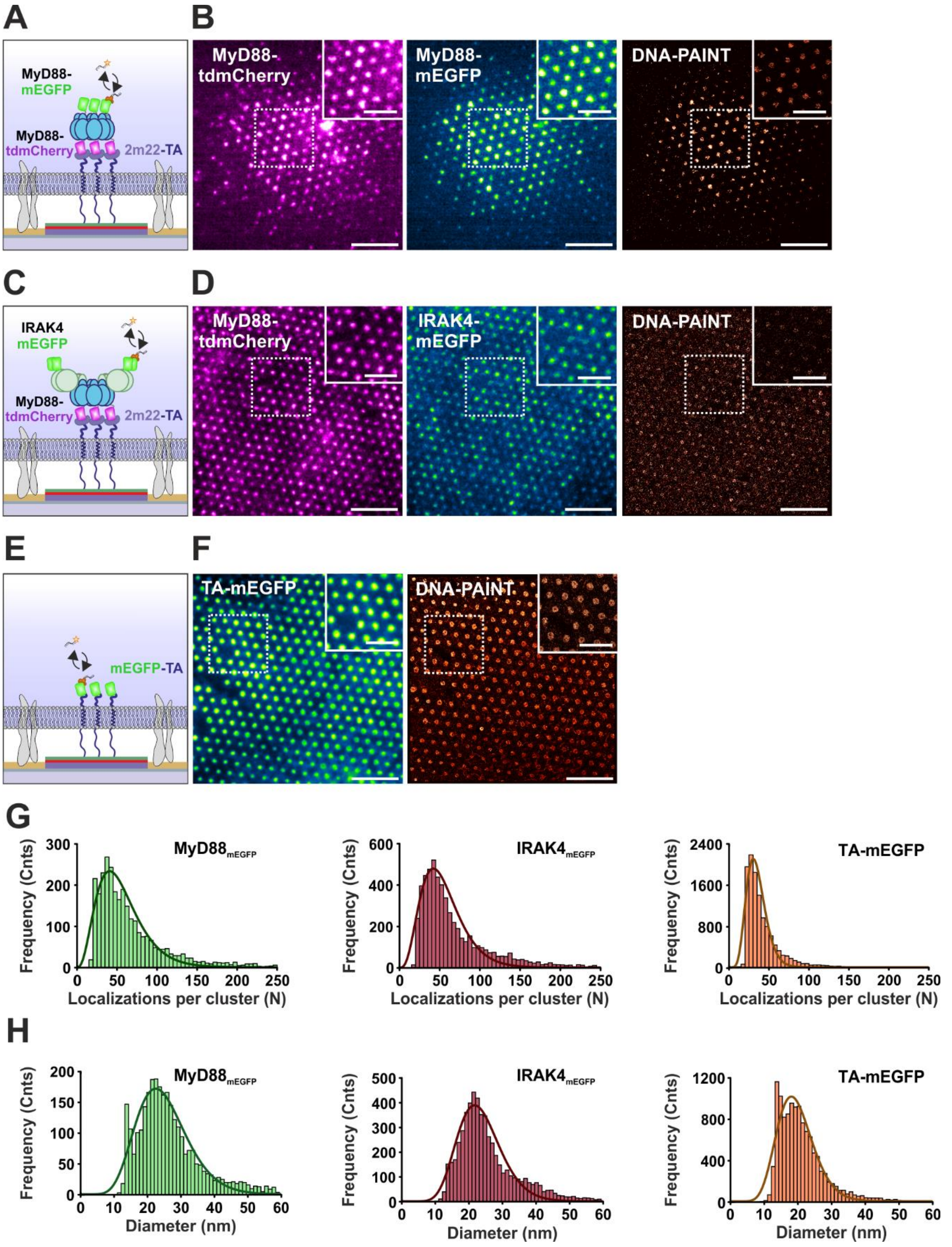


**Figure S10. Comparative DNA-PAINT analysis of clustering characteristics for TA-mEGFP relative to MyD88-mEGFP and IRAK4-mEGFP.** (A, C, E) Schematic representation of the experimental setup for DNA-PAINT imaging of MyD88-mEGFP (A), IRAK4-mEGFP (C), and TA-mEGFP (E). (B, D, F) Representative dual-color TIRF microscopy images of immobilized proteins: MyD88-tdmCherry with co-recruited MyD88-mEGFP (B), MyD88-

tdmCherry with co-recruited IRAK4-mEGFP (D), and TA-mEGFP (F). Corresponding DNA-PAINT super-resolution images highlight protein localization patterns within individual nanodots Insets show magnified views of individual nanodots. Scale bars: 5 µm; insets: 2 µm. (G, H) Quantitative distribution analyses of localizations per cluster (H) and cluster diameter (I) for MyD88-mEGFP (green), IRAK4-mEGFP (red), and TA-mEGFP (orange). Histograms fitted with gamma distributions.

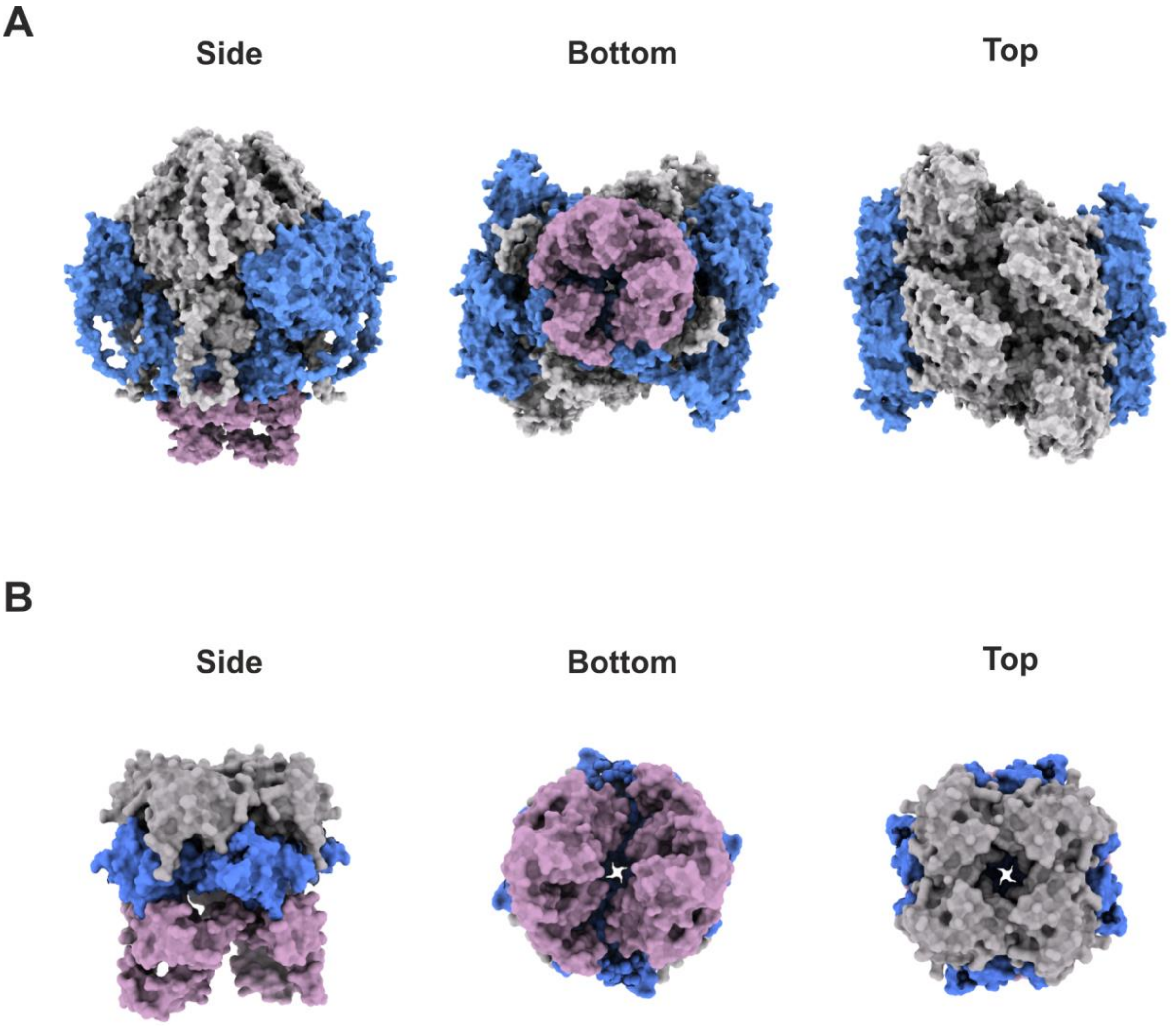


**Figure S11. Structural modeling of the myddosome complex using AlphaFold3.** (A) AlphaFold3-derived structural model of the complete myddosome assembly, illustrating the oligomeric and hierarchical architecture with domain-specific color coding: 6x hMyD88 (M54-I109) death domain (DD) in light pink, 4x mIRAK4 (M1-A460) in blue, and 4x mIRAK1 (M1-F688) in grey. The side (left), bottom (middle), and top (right) views emphasize the spatial arrangement of individual components, revealing intricate inter-domain interactions crucial for complex integrity. (B) Refined structural representation focusing exclusively on the DD of hMyD88 (M54-I109, light pink), mIRAK4 (R20-A104, blue), and mIRAK1 (M27-A106, grey). The side (left), bottom (middle), and top (right) views show the symmetric oligomerization and bottom-up assembly of the complex.

### Supplementary Tables

#### Table S1. Recombinant proteins used for *in vitro* experiments

| Protein/Denomination | Plasmid name | Source |
|---|---|---|
| Halo-mEGFP[1] | pET-21a HaloTag-mEGFP-H6 | Philippi, *et al*., 2022 |
| SpyCatcher-sfGFP[1] | pDEST14 H6-TEV-SpyCatcher003-sfGFP | Addgene (#133449) |
| ALFAnb-mEGFP[1] | pET-21a H6-aALFAnb-mEGFP | This manuscript |
| Halo-tdR7[1] | pET-21a DARPinR7-linker[2]-HaloTag-DARPinR7-H8 | Philippi, *et al*., 2022 |
| SpyCatcher-tdR7[1] | pET-21a DARPinR7-linker[2]-SpyCatcher003-DARPinR7-H8 | This manuscript |
| ALFAnb-tdR7[1] | pET-21a DARPinR7-linker[2]-aALFAnb-DARPinR7-H8 | This manuscript |
| ALFAnb-td2m22[1] | pET-21a DARPin2m22-linker[2]-aALFAnb-linker[2]-DARPin2m22-H6 | This manuscript |

1) Protein concentration: 1 µM; Incubation time: 30 min, 1 h for HaloTag recombinant proteins.
2) Linker: $(GSGSD)_3$

#### Table S2. Description of plasmids used for live cell cNDA experiments

| Plasmid name | Denomination | Source |
|---|---|---|
| pSems-leader-ALFAnb-GSlinker-TMD[1]-GSlinker-GFPnb | TA-GFPnb | Philippi, *et al*., 2022 |
| pSems-leader-ALFAnb-GSlinker-TMD[1]-GSlinker-2m22 | TA-2m22 | This manuscript |
| pSems-leader-Spycatcher003-GSlinker-TMD[1]-GSlinker-GFPnb | TS-GFPnb | This manuscript |
| pSems-leader-ALFAnb-GSlinker-TMD[1]-mEGFP | TA-mEGFP | This manuscript |
| pSems-mEGFP | mEGFP | Löchte, *et al*., 2014 |
| pSems-tdmCherry | tdmCherry | This manuscript |
| pSems-hMyD88-tdmCherry | MyD88-tdmCherry | This manuscript |
| pSems-hMyD88-mEGFP | MyD88-mEGFP | This manuscript |
| pSems-mIRAK4-mEGFP | IRAK4-mEGFP | This manuscript |
| pSems-mIRAK4-mtagBFP | IRAK4-mtagBFP | This manuscript |

| | | |
|---|---|---|
| pSems-mIRAK1-mEGFP | IRAK1-mEGFP | This manuscript |
| pSems-tdiRFP-hTRAF6 | TRAF6-tdiRFP | This manuscript |

[1] Transmembrane domain (TMD): Artificial TMD with the sequence of $(ALA)_7$ repeats.

**Table S3. List of key materials and suppliers**

| **Reagent and abbreviation** | **Source** | **Catalogue number** |
|---|---|---|
| Glass coverslips | Marienfeld Laboratory glassware | 0117640 |
| (1-Naphthylmethyl)trichlorosilane (NMTS) | Gelest | SIN6596.0 |
| (4-chlorophenyl)triethoxysilane (CPTES) | Sigma Aldrich | 597910 |
| Vinyltrimethoxysilane (VTMS) | Sigma Aldrich | 235768 |
| Toluene | Fisher Scientific | 14214914 |
| Poly-L-lysine -hydrochloride (PLL) | Sigma Aldrich | 2658 |
| α-Butyric Acid NHS Ester-ω-Maleimidohexanamido $PEG_{3k}$ | Rapp Polymere | 133000-65-35 |
| $HTL-NH_2$ | Own synthesis | Liße, *et al*., 2011 |
| Albumin Fraction V from bovine serum (BSA) | Carl Roth | 0136.1 |
| Peptides (ALFAtag, SpyTag) | Romer Labs | Custom synthesis |
| N-Succinimidyl 13-Maleimido-11-oxo-4,7-dioxa-10-azatridecanoate (Mal-$PEG_2$-NHS) | Tokyo Chemical Industry | M3079 |
| N-2-Hydroxyethylpiperazine-N'-2-ethane sulphonic acid (HEPES) | Carl Roth | 6763.3 |
| Protease inhibitor | Serva | 39106 |
| DNase | Sigma Aldrich | DN25 |
| Lysozyme | Sigma Aldrich | L6876 |
| Isopropyl-β-D-thiogalactopyranoside (IPTG) | Thermo Fisher Scientific | R0392 |
| Imidazole | Carl Roth | 3899.4 |
| Urea | Carl Roth | 3957.1 |
| Ethylenediamine tetra-acetic acid (EDTA) | Carl Roth | 8040.215 |
| MASSIVE-TAG-X2-FAST anti-GFP | Massive Photonics | FAST-DNA-PAINT Kit |
| MASSIVE-RESI anti-GFP | Massive Photonics | RESI Kit |
| 90 nm standard gold | Cytodiagnostics | G-90-20 |

| | | |
|---|---|---|
| Nanoparticles | | |
| Dulbecco's phosphate buffered saline (PBS) | PanBiotech | P04-36500 |
| Dulbecco's Modified Eagle's Medium (MEM) | PanBiotech | P04-09500 |
| Fetal bovine serum (FBS) | PanBiotech | P30-3031 |
| Trypsin | Capricorn Scientific | TRY-1B |
| Linear polyethylenimine hydrochloride (PEI) | Polysciences | 24765-1 |
| NaCl | Carl Roth | 3957.1 |

**Table S4. Quantitative characterization of bNDAs obtained from *in vitro* DNA-PAINT experiments**

| Condition | Diameter (nm) | Peak intensity (Cnts) | Loc. Density (per 1000 nm$^2$) | Contrast |
|---|---|---|---|---|
| CPTES_PPA GFPnb | 550 ± 25 | 7599 ± 766 | 28 ± 3 | 30 ± 8 |
| VTMS_PPA GFPnb | 457 ± 35 | 12369 ± 1488 | 50 ± 7 | 66 ± 34 |
| VTMS_BSA GFPnb | 530 ± 24 | 11491 ± 1931 | 38 ± 5 | 54 ± 24 |
| CPTES_PPA ALFAnb | 551 ± 26 | 5515 ± 748 | 19 ± 2 | 20 ± 4 |
| VTMS_PPA ALFAnb | 374 ± 27 | 16533 ± 2684 | 83 ± 13 | 152 ± 47 |
| VTMS_BSA-A ALFAnb | 540 ± 17 | 20898 ± 2565 | 65 ± 6 | 100 ± 32 |

**Table S5. Summary of cNDA cluster analysis obtained from DNA-PAINT experiments**

| Target | Clustered localizations per cNDA [%] | Localizations per cluster [N] | Cluster diameter [nm] | Cluster area [nm$^2$] |
|---|---|---|---|---|
| MyD88-mEGFP | 57 ± 10 | 41 ± 19 | 22 ± 6 | 233 ± 145 |
| IRAK4-mEGFP | 73 ± 6 | 42 ± 17 | 22 ± 5 | 216 ± 111 |
| TA-mEGFP | 32 ± 13 | 30 ± 10 | 18 ± 4 | 142 ± 71 |

**Table S6. Quantitative RESI analysis of IRAK4-mEGFP in cNDAs**

| | IRAK4-mEGFP |
|---|---|
| **Number of cNDAs [N]** | 378 |
| **Number of molecules [N]** | 9691 |
| **Localization precision [Å]** | 5.6 ± 1.6 |
| **Number of molecules per cNDA [N]** | 20 ± 7 |
| **RESI localization density [$\mu m^{-2}$]** | 73 ± 27 |

**Table S7. Distribution of oligomeric states and spatial organization of IRAK4-mEGFP molecules determined by RESI**

| Class | Count [N] | Percent [%] | Intermolecular distance [nm] |
|---|---|---|---|
| Monomer | 5159 | 53 | N.D. |
| Dimer | 2794 | 29 | 12 ± 5 |
| Trimer | 1155 | 12 | 15 ± 7 |
| Tetramer | 308 | 3 | 18 ± 8 |
| Higher Oligomer | 275 | 3 | N.D. |
